# One-carbon metabolism nutrients impact the interplay between DNA methylation and gene expression in liver, enhancing protein synthesis in Atlantic Salmon

**DOI:** 10.1101/2023.09.18.558254

**Authors:** Takaya Saito, Marit Espe, Vibeke Vikeså, Christoph Bock, Tårn H. Thomsen, Anne-Catrin Adam, Jorge M.O. Fernandes, Kaja H. Skjaerven

## Abstract

Supplementation of one-carbon (1C) metabolism micronutrients, which include B-vitamins and methionine, is essential for the healthy growth and development of Atlantic salmon (*Salmo salar*). However, the recent shift towards non-fish meal diets in salmon aquaculture has led to the need for reassessments of recommended micronutrient levels. Despite the importance of 1C metabolism in growth performance and various cellular regulations, the molecular mechanisms affected by these dietary alterations are less understood.

To investigate the molecular effect of 1C nutrients, we analysed gene expression and DNA methylation using two types of omics data: RNA sequencing (RNA-seq) and reduced-representation bisulfite sequencing (RRBS). We collected liver samples at the end of a feeding trial that lasted 220 days through the smoltification stage, where fish were fed three different levels of four key 1C nutrients: methionine, vitamin B6, B9, and B12.

Our results indicate that the dosage of 1C nutrients significantly impacts genetic and epigenetic regulations in the liver of Atlantic salmon, particularly in biological pathways related to protein synthesis. The interplay between DNA methylation and gene expression in these pathways may play an important role in the mechanisms underlying growth performance affected by 1C metabolism.

**Author Summary:** Atlantic salmon rely on one-carbon (1C) metabolism micronutrients like B-vitamins and methionine, which they acquire through their diets. Small pelagic fish are the primary source in the wild, but finding sustainable alternatives such as plants, insects, and algae is challenging as salmon aquaculture expands. Adjusting nutrient levels when changing base ingredients further complicates the task. Understanding the molecular mechanisms affected by these micronutrients is crucial for aquaculture sustainability.

In this study, we investigated the molecular effects of 1C metabolism micronutrients on Atlantic salmon. Liver samples from salmon fed varying levels of key 1C nutrients over a 220-day trial were analysed using RNA sequencing (RNA-seq) and reduced-representation bisulfite sequencing (RRBS) to assess gene expression and DNA methylation, respectively.

Our results revealed significant impacts of 1C nutrient dosage on genetic and epigenetic regulations in the salmon liver, particularly in protein synthesis pathways. The interplay between DNA methylation and gene expression in these pathways influences growth performance under 1C metabolism.

Uncovering molecular changes resulting from dietary alterations provides valuable insights to optimize nutritional requirements in salmon aquaculture, supporting sustainable production and welfare of this important species.

## Introduction

One-carbon metabolism micronutrients, or simply 1C nutrients, are essential micronutrients for healthy animal development [1], including the development over the smoltification stage of Atlantic salmon (*Salmo salar*) [2]. While three B-vitamins, pyridoxine (vitamin B6), folate (vitamin B9), and cobalamin (vitamin B12), play a crucial role in maintaining 1C metabolism [3, 4], fish need to acquire them through their diets as they cannot produce folate and vitamin B12 on their own. The main source of these vitamins for wild salmon is small pelagic fish, but the rapid development of plant-based diets in the salmon aquaculture industry has prompted reassessments of recommended levels of micronutrients in their diets [5, 6]. Nonetheless, it remains challenging to determine the optimal range of 1C nutrients, such as B-vitamins and methionine, that support healthy and robust fish growth across different conditions and life stages. Moreover, there is an urgent need to explore alternative protein sources that do not rely on fishmeal or plant-based diets to enhance the sustainability of salmon aquaculture.

Methionine is a key component in 1C metabolism, providing the methyl group required for various biological functions, such as DNA methylation [7], post-translational protein modifications [8], creatine synthesis [9], carnitine synthesis [10], endogenous choline synthesis [11, 12], and polyamine synthesis [13]. Before methionine can function as a methyl donor, it needs to be converted to S-adenosylmethionine (SAM) by the enzyme methionine adenosyl transferase (MAT) [14]. After donating the methyl group, SAM is converted to S-adenosylhomocysteine (SAH). As SAH is unstable and easily converted to homocysteine, which is cytotoxic to the cells [15], it needs to be either transsulfurated or remethylated to methionine. 1C metabolism is closely related to the methionine cycle and the folate cycle as they transfer and utilize one-carbon units within cells. The former involves methionine, SAM, SAH, and vitamin B12, while the latter involves vitamin B6, folate, and B12. Vitamin B6 is necessary for transsulfuration, while folate and B12 are used for the remethylation of homocysteine [16].

The liver is a key target organ for 1C metabolism and methylation reactions. Plant-based diets were found to increase lipid accumulation in the liver and intestinal tissues [17], and sub-optimal methionine level in the diets affected hepatic triglycerides (TAGs) accumulation in Atlantic salmon [18]. In addition, salmon fed low B-vitamin diets grew fattier than those fed higher B-vitamin diets [6]. Liver lipid accumulation is associated with increased metabolic stress, energy depletion, cytokine activation, and inflammation in mammals [19-21].

In the present study, we used liver samples from Atlantic salmon fed three different dietary levels of 1C nutrients, including methionine, vitamin B6, folate, and vitamin B12. Previous findings from this feeding trial showed that a medium dosage of the 1C nutrients improved growth and reduced liver size, while a high dosage had limited contribution [2]. To gain insights into the molecular mechanisms underlying these effects, we investigated two types of omics data, namely RNA-seq for gene expression and reduced-representation bisulfite sequencing (RRBS) for DNA methylation. Our goal was to gain a better understanding of the affected biological functions and identify the interactions between them by integrating these datasets.

## Results

### Atlantic salmon fed with varied 1C nutrient levels showed differential growth performance

The present study examines the impact of varied 1C nutrient levels on the gene expression and DNA methylation patterns in the liver of Atlantic salmon. The feeding trial lasted 220 days through the smoltification stage (Fig 1A), and three diets were used: Ctrl, 1C+, and 1C++, each containing different levels of four key 1C nutrients: methionine, folate, vitamin B6, and vitamin B12 (Fig 1B). While the Ctrl group was fed with the recommended levels of 1C nutrients, the 1C+ and 1C++ groups were given higher levels of these nutrients (Fig 1C & Table S1 in S1 Info). The growth performance and nutrient status of the fish were previously assessed, and the results have been summarized and discussed elsewhere [2].

**Fig 1.**
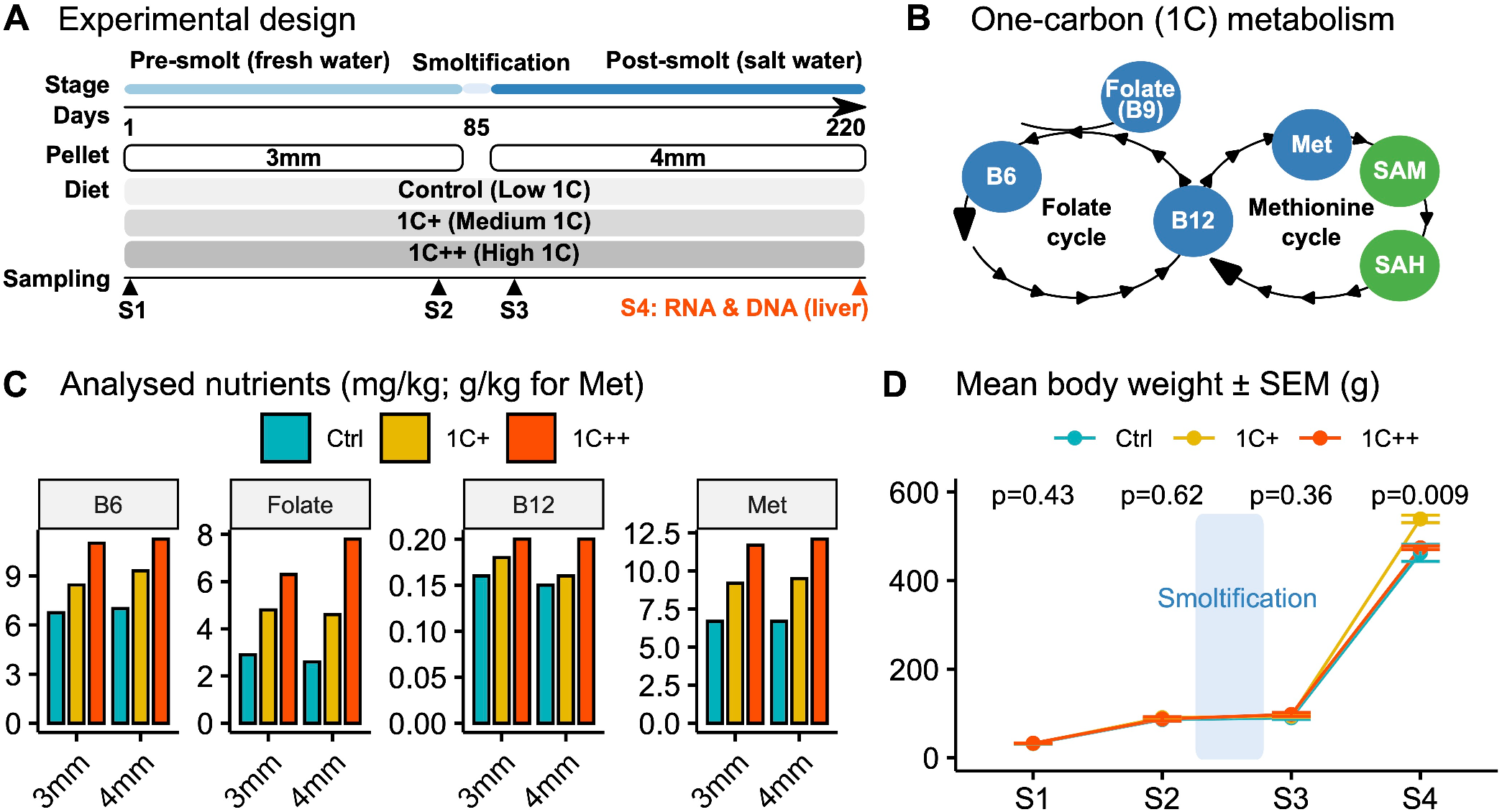
Overview of experimental design and nutrient levels in the feeding trial with growth performance. **(A)** A schematic diagram depicting different stages, pallet sizes, diets, and sampling points throughout the trial. **(B)** Diagram showing targeted nutrients (blue) along with SAM and SAH (green) in the 1C metabolism that includes folate and methionine cycles. **(C)** Barplots showing levels of four 1C nutrients (B6, folate, B12, and methionine) analysed for both 3mm and 4mm pellets for three diet groups (Ctrl, 1C+, and 1C++). **(D)** Line plot showing the growth rates in body weights (g) for three diet groups along with p-values calculated by ANOVA.

At the final sampling point (S4), the fish in the 1C+ group had a significantly better growth performance than those in the other two groups, with an average weight that was ∼50 grams greater (Fig 1D). Other measures, including condition factor and hepatosomatic index, also indicated a healthier and better growth for the 1C+ group. In contrast, the growth performance of the 1C++ group was only slightly better than that of the Ctrl group (Table S2 in S1 Info).

To uncover the underlying genomic and epigenetic regulations associated with the observed differences in growth performance over smoltification, we analysed gene expression and DNA methylation of liver samples collected at the S4 sampling point (post-smolt stage). The remaining sections of the Results will focus on the analysis of these data.

### Clustering analysis revealed two strong gene expression patterns that are common to the 1C-supplemented groups

We conducted an RNA-seq analysis on 27 liver samples from three groups (n=9) to investigate the effect of 1C nutrient levels on gene expression. We identified 22,066 expressed genes by aligning read counts to the reference genome (detailed statistics for each sample are provided in Table S3 in S1 Info). Principal component analysis (PCA) showed a clear separation between the 1C-supplemented groups (1C+ and 1C++) and the control group (Fig 2A).

**Fig 2.**
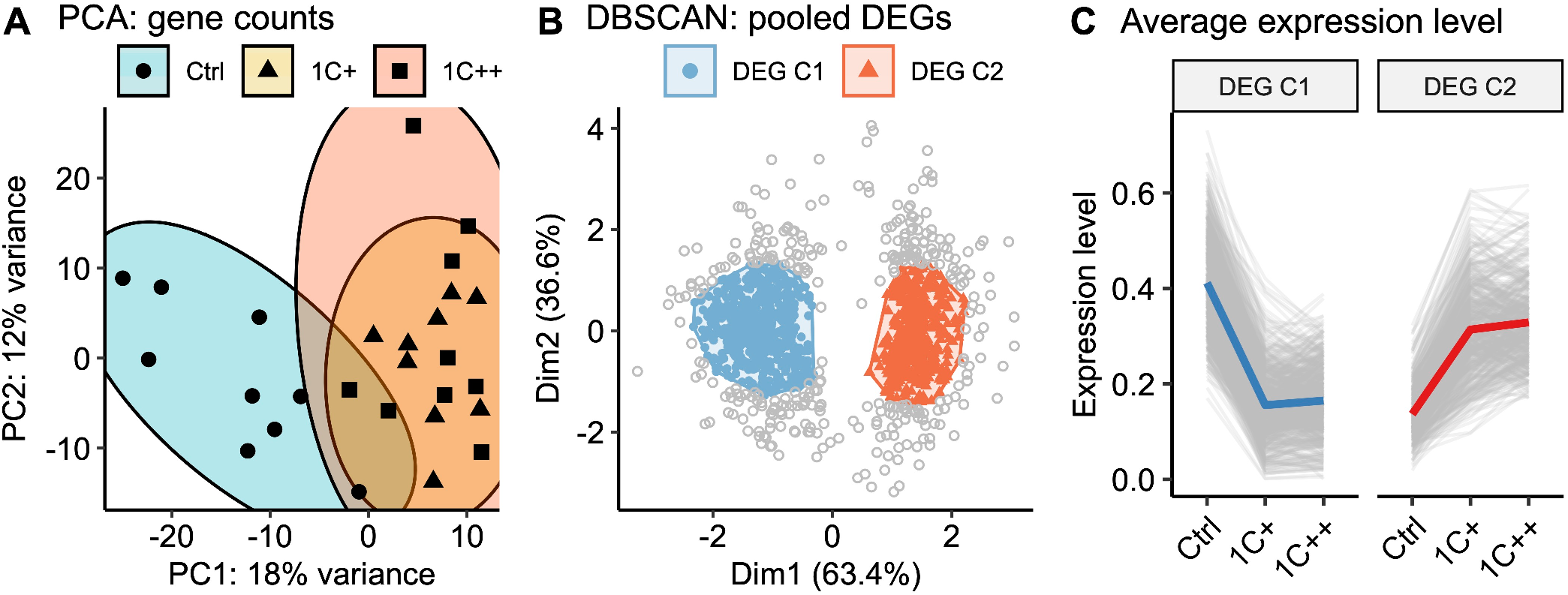
Clustering analysis of gene expression differences among three diets. **(A)** A PCA plot displaying the clusters of three diet groups - Ctrl (blue, semi-transparent), 1C+ (yellow, semi-transparent), and 1C++ (red, semi-transparent) - using 27 RNA-seq samples including nine samples from each group. Top 1000 high variance genes were used as input data. (B) A dot plot showing the DBSCAN result in a PCA format with two identified clusters, DEG C1 (blue) and DEG C2 (red), on the pooled set of DEGs generated from three pair-wise comparisons (1C+ vs Ctrl, 1C++ vs Ctrl, 1C++ vs 1C+). (C) Line plots showing normalised read counts as expression levels for three diet groups with the total averages for DEG C1 (blue) and DEG C2 (red) clusters.

Subsequently, we created a pooled set of differentially expressed genes (DEGs) by selecting genes that were differentially expressed in at least one of the three pair-wise comparisons (1C+ vs Ctrl, 1C++ vs Ctrl, and 1C++ vs 1C+; counts of DEGs from each comparison were provided in Table S4 in S1 Info). This pooled set allowed us to directly use normalised read counts as gene expression level instead of log fold changes, which is useful to identify interesting expression patterns, for example by using a robust clustering method, such as density-based spatial clustering of applications with noise (DBSCAN).

The DBSCAN clustering on the 1268 pooled DEGs showed that 74% (939 DEGs) of them were linked to two major clusters (DEG C1 and DEG C2) consisting of 533 and 409 DEGs, respectively (Fig 2B). The gene expression patterns of these clusters were concordant and demonstrated that all the genes in DEG C1 were down-regulated, while all the genes in DEG C2 were up-regulated in the 1C-supplemented groups (1C+ and 1C++) compared to the control group (Fig 2C).

The clustering analysis strongly suggests that both medium and high levels of 1C nutrients (1C+ and 1C++, respectively) affected gene expression profiles in a similar manner in the liver of Atlantic salmon, despite the significant difference in growth performance between them.

### Functional analysis identified 14 biological pathways significantly affected by 1C nutrient supplementation

To examine the impact of 1C nutrient supplementation on biological pathways, we conducted functional analysis using the Kyoto Encyclopedia of Genes and Genomes (KEGG) database by employing two methods, namely over-representation analysis (ORA) and gene set enrichment analysis (GSEA). ORA was performed on DEGs, while GSEA was conducted on the whole gene set (see Materials and Methods for details). In total, ORA and GSEA together identified 14 KEGG pathways that can be grouped into three categories: metabolism, genetic information processing, and cellular processes (Table 1; detailed lists are provided in Table S5 in S1 Info for ORA and Tables S6-S8 in S1 Info for GSEA).

**Table 1.** Enriched KEGG pathways by ORA and GSEA.

| Category | Sub-category | KEGG pathway | ORA<sup>1</sup> | GSEA<sup>2</sup> |
| --- | --- | --- | --- | --- |
| Metabolism | Global and overview maps | Biosynthesis of cofactors | Down | Down (1C++) |
|  |  | Biosynthesis of amino acids | Down | Down |
|  | Energy metabolism | Oxidative phosphorylation | - | Up |
|  | Lipid metabolism | Steroid biosynthesis | - | Down |
|  |  | Linoleic acid metabolism | Up | Up (1C+) |
|  | Amino acid metabolism | Cysteine and methionine metabolism | Down | Down (1C++) |
|  |  | Arginine biosynthesis | Down | - |
|  | Xenobiotics biodegradation and metabolism | Drug metabolism - cytochrome P450 | Down | - |
|  |  | Metabolism of xenobiotics by cytochrome P450 | Up | - |
| Genetic Information Processing | Translation | Ribosome | - | Up |
|  |  | Aminoacyl-tRNA biosynthesis | - | Down (1C++) |
|  | Folding, sorting and degradation | Protein processing in endoplasmic reticulum | Up | Up (1C+) |
|  |  | Protein export | Up | Up (1C+) |
| Cellular Processes | Cell growth and death | Cell cycle | - | Down |

<sup>1</sup>Down: enriched in C1 cluster, Up: enriched in C2 cluster, and "-": not enriched by ORA. <sup>2</sup>Down and Up: enriched by both 1C+ and 1C++ groups with negative and positive NESs, respectively, but enriched only by one group when a group name (1C+ or 1C++) is followed. "-": not enriched by GSEA.

All the six pathways supported by both ORA and GSEA showed concordant regulations regarding up- and down-regulation. Specifically, all the four pathways related to amino acid metabolism were down-regulated, whereas the remaining two pathways related to protein processing were up-regulated. Moreover, several pathways showed opposite regulations even though they were in the same subcategories. For instance, steroid biosynthesis was down-regulated, while linoleic acid metabolism was up-regulated, and both pathways are associated with lipid metabolism. In addition, two pathways related to cytochrome P450 also showed both up- and down-regulation (Table 1).

These results suggest that supplementation of both medium and high levels of 1C nutrients (1C+ and 1C++) potentially bypassed part of amino acid metabolism and enhanced protein processing in the liver of Atlantic salmon. Moreover, several pathways showed both up- and down-regulation, even though they belonged to the same subcategories, specifically in metabolism related pathways inducing lipid metabolism and xenobiotics biodegradation by cytochrome P450.

### Supplementation of 1C nutrients strongly influenced expression of genes encoding enzymes

To identify genes strongly affected by 1C nutrients, we filtered out DEGs with two threshold values of log fold changes (LFCs), namely stringent (|LFC| > 2) and semi-stringent (|LFC| > 1.5) thresholds (see Materials and Methods for details). This approach resulted in 10 and 62 genes for stringent and semi-stringent thresholds, respectively.

Among the 10 genes identified by the stringent threshold, seven belonged to the DEG C1 cluster, indicating strong down-regulation by the 1C-supplemented groups (1C+ and 1C++) compared to the control group (Table 2). Moreover, all of them were enzyme-coding genes except for two uncharacterised genes. Specifically, two paralogs of the *mat2* gene, which is a rate-limiting enzyme for SAM synthesis [14], were down-regulated in a dose-dependent manner, with the 1C++ group showing more down-regulation than the 1C+ group when compared to the control group. Additionally, two paralogs of the *lnx* gene, which assists in the transfer of ubiquitin to target proteins [22], were also down-regulated, but the 1C+ group was more down-regulated compared to the 1C++ group.

**Table 2.**
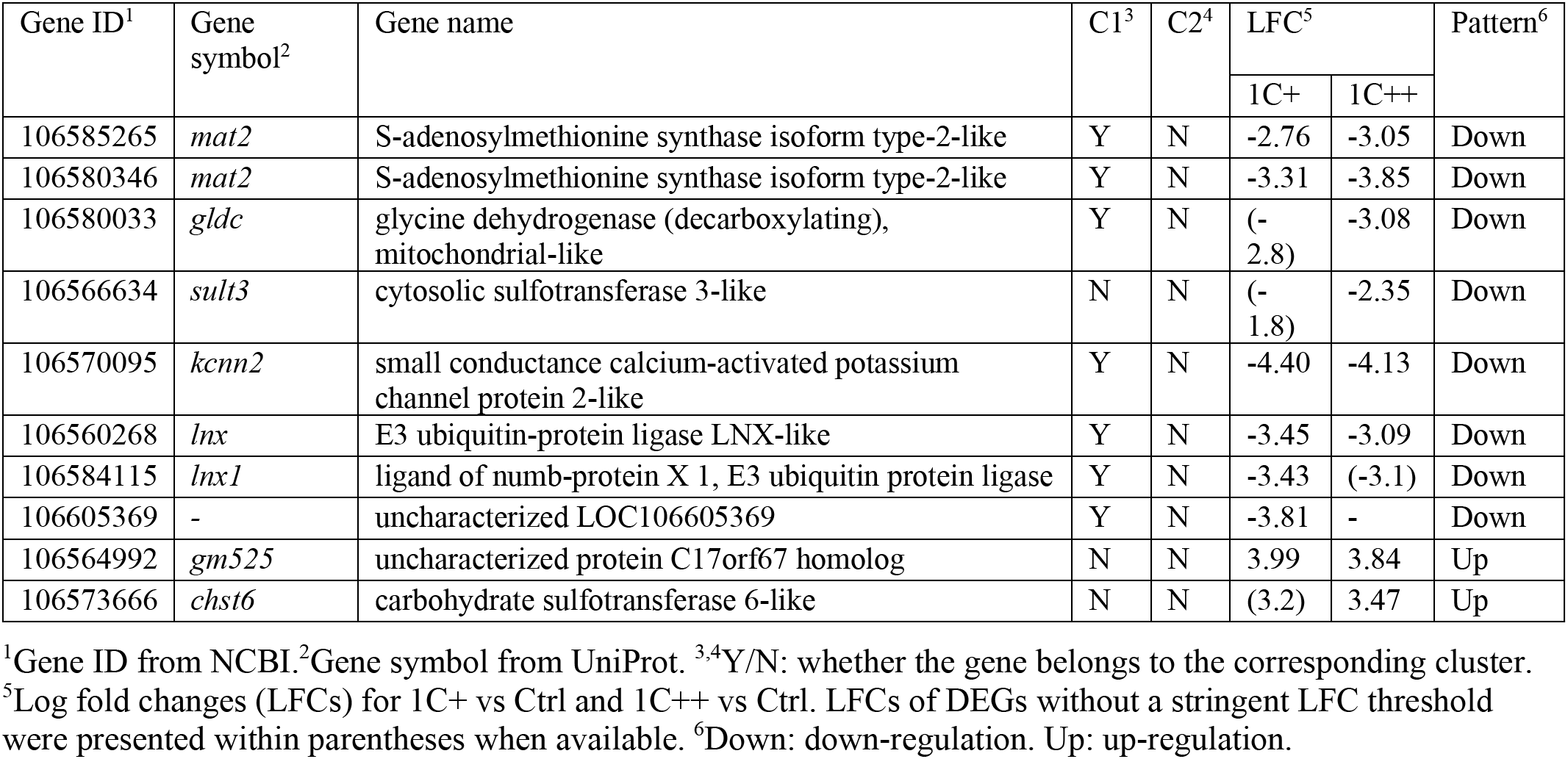
List of DEGs with a stringent LFC threshold (|LFC| > 2).

As the stringent threshold resulted in no DEGs from the 1C++ vs 1C+ comparison, we also analysed the results of the semi-stringent threshold to identify genes that were potentially responsible for the limited growth performance of the 1C++ group. Among the 62 genes identified by the semi-stringent threshold, only one gene, *cyp27b1* (gene ID: 106583283, gene name: 25-hydroxyvitamin D-1 alpha hydroxylase, mitochondrial-like), was from the 1C++ vs 1C+ comparison. The *cyp27b1* gene, which encodes a cytochrome P450 enzyme [23], exhibited the lowest expression level for the 1C++ group in an order of 1C++ < Ctrl < 1C+.

To summarise, 1C nutrient supplementation strongly influenced the regulation of the genes that encode various enzymes, including SAM synthesis ubiquitination, and cytochrome P450. Moreover, altered gene expression of the *mat2* gene affected the regulation of SAM synthesis, which potentially influenced DNA methylation profiles.

### The SAM/SAH ratio indicated a low methylation capacity in fish given a high dosage of 1C nutrients

The SAM/SAH ratio is commonly used as marker to assess methylation capacity [24], and both SAM and SAH play a crucial role in the methionine cycle (Fig 1B). We performed reversed-phase chromatography to measure SAM and SAH levels in three feeding groups in the liver of Atlantic salmon and found that the SAM level was significantly lower in the 1C++ group compared to the other two groups. Although SAH level was higher in the 1C++ group, the difference was not statistically significant. The SAM/SAH ratio for the 1C++ group was significantly lower than that for the other groups, indicating lower methylation capacity for the 1C++ group (Fig 3A).

**Fig 3.**
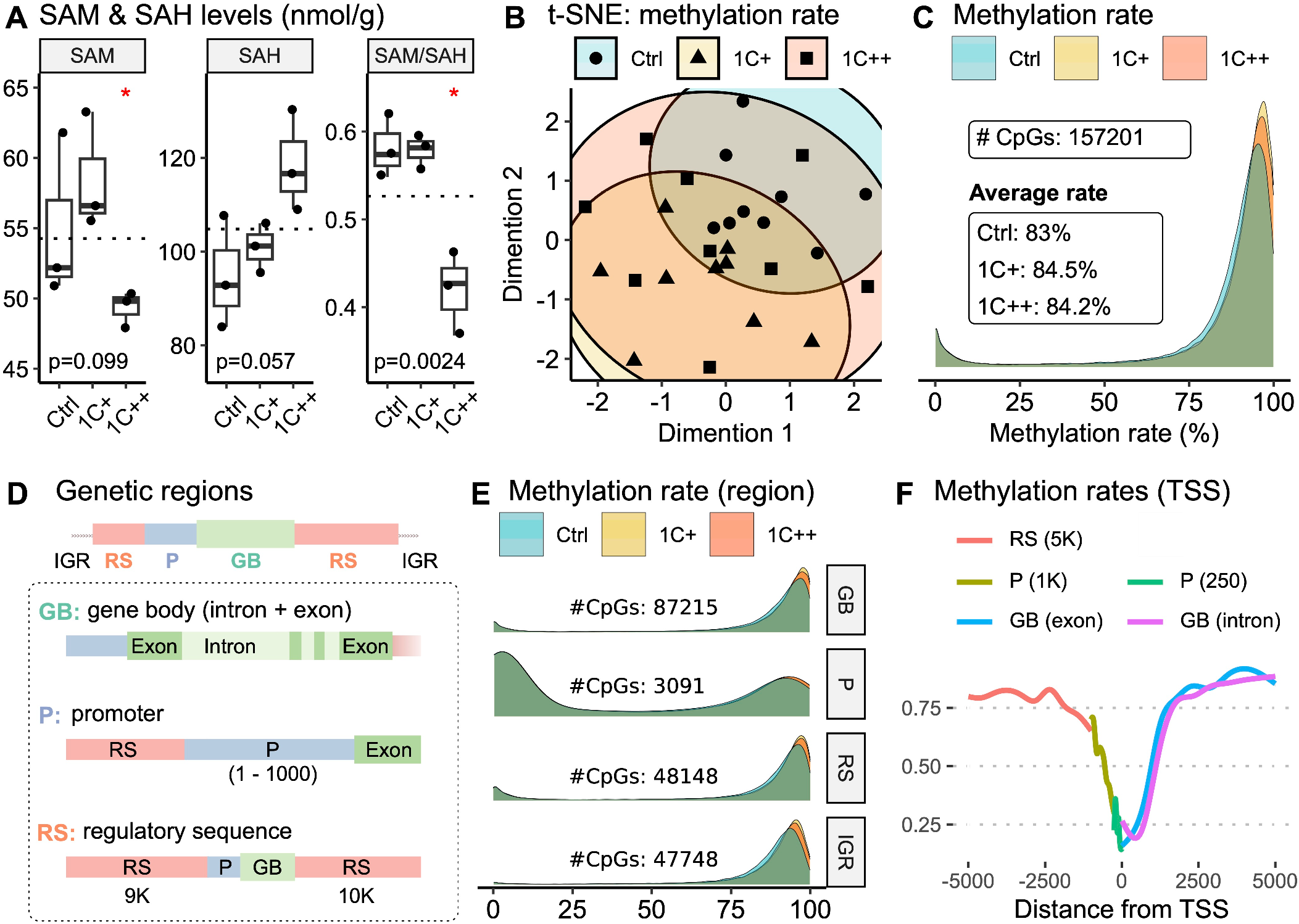
Global and regional DNA methylation landscapes along with SAM/SAH ratio. **(A)** Box plots showing SAM and SAH levels (nmol/g) with SAM/SAH ratio for three diet groups. P-values are calculated by ANOVA, and red stars on the top indicate statistical significance from the base mean by t-test. Dotted horizontal lines indicate base means. **(B)** A t-SNE plot displaying the clusters of three diet groups - Ctrl (blue, semi-transparent), 1C+ (yellow, semi-transparent), and 1C++ (red, semi-transparent) - using 27 RRBS samples. **(C)** A ridge density plot showing average methylation rates of 157,201 CpG sites for three diet groups. **(D)** A schematic diagram showing the definition of three different genomic regions - gene body (GB, green), promoter (P, blue), and regulatory sequence (RS, red) along with intergenic regions (IGRs). **(E)** Ridge density plots showing average methylation rates for three diet groups in four different genetic regions. **(F)** A running average line plot showing average DNA methylation rates of all RRBS samples within 5000 bp up- and down-stream around TSSs. The running average lines were calculated separately for five different genetic regions - RS (red), P (dark yellow and green), and GB (blue and pink).

These findings suggest that a high dosage of 1C nutrients (1C++) may impair methylation capacity, which can have implications for the lower growth performance of the 1C++ group compared to the 1C+ group.

### Supplementation of 1C nutrients broadly contributed to the shifting of DNA methylation rates that are positively correlated with growth performance

We conducted an RRBS analysis on liver samples from three groups (n=9) to examine the effect of 1C nutrient levels on DNA methylation profiles. After aligning the sequence reads to the salmon genome (see Table S9 in S1 Info for alignment statistics), we identified over 150,000 CpG sites that were methylated.

Despite using various clustering approaches, including t-distributed stochastic neighbour embedding (t-SNE), we found no clear clustering of the CpG sites among the three feeding groups (Fig 3B for t-SNE and Fig S1 in S1 Info for other clustering results). This implies that the individual variability in methylation profiles was higher than the group variability.

However, we observed noticeable peak shifts in methylation rates (Fig 3C), with the distributions of methylation rates being significantly shifted in an order of Ctrl < 1C++ < 1C+ (Kolmogorov–Smirnov test; Table S10 in S1 Info). This shift was consistent with the observed growth performance (e.g., mean body weight shown in Fig 1D). Additionally, we found that the shift was more pronounced for CpG sites with high methylation rates (Fig 3C, between 75% and 100%), indicating that these highly methylated sites were even more methylated in the 1C-supplemented groups (1C+ and 1C++) compared to the control group (Fig 3C).

### Shifting patterns of DNA methylation rates were similar in different genomic regions

To investigate region-specific methylation patterns, we annotated CpG sites into four genomic regions (Fig 3D): gene body (GB), promoter (P), regulatory sequence (RS), and intergenic region (IGR). Similar to the shift that we observed in the whole genome region, the distributions of methylation rates were significantly shifted in an order of Ctrl < 1C++ < 1C+ in all four genetic regions, except for the comparison between 1C+ and 1C++ in the promoter region that showed no significant difference (Fig 3E and Table S10 in S1 Info).

The underlying methylation rates appeared to be different in the promoter regions compared to the other regions, as more than half of the CpG sites were associated with low methylation rates (Fig 3E, between 0% and 25%). This can be explained by low methylation rates around transcription start sites (TSSs). We observed that methylation rates decreased sharply to around 25%, with the lowest point at the transcription start sites (TSSs), from an average methylation rate of approximately 80% in the surrounding area (Fig 3F).

Our findings suggest that 1C nutrient supplementation modulated methylation profiles similarly in different regions but with an exception in the regions around TSSs, which showed low methylation rates and less shifting.

### Differential methylation analysis revealed more hyper-methylated CpGs in the 1C-supplemented groups when compared to the control group

We performed a differential methylation analysis to identify CpG sites that were differentially methylated with statistical significance between treatment groups. We used three pair-wise comparisons (1C+ vs Ctrl, 1C++ vs Ctrl, and 1C++ vs 1C+), which lead to over 6000 differentially methylated CpG sites (DMCs) for each comparison (Table S11 in S1 Info).

The result showed that approximately 70% of DMCs identified in two comparisons (1C+ vs Ctrl and 1C++ vs Ctrl) were hyper-methylated (Fig 4A). Additionally, the 1C+ group had slightly more hyper-methylated DMCs than the 1C++ group (shown as hypo-methylated DMCs in Fig 4A as the 1C+ group was used as a reference/base group). This finding is consistent with our observation that methylation rates were shifted in the order of Ctrl < 1C++ < 1C+ with the mean methylation rates of 83%, 84.2%, and 84.5%, accordingly.

**Fig 4.**
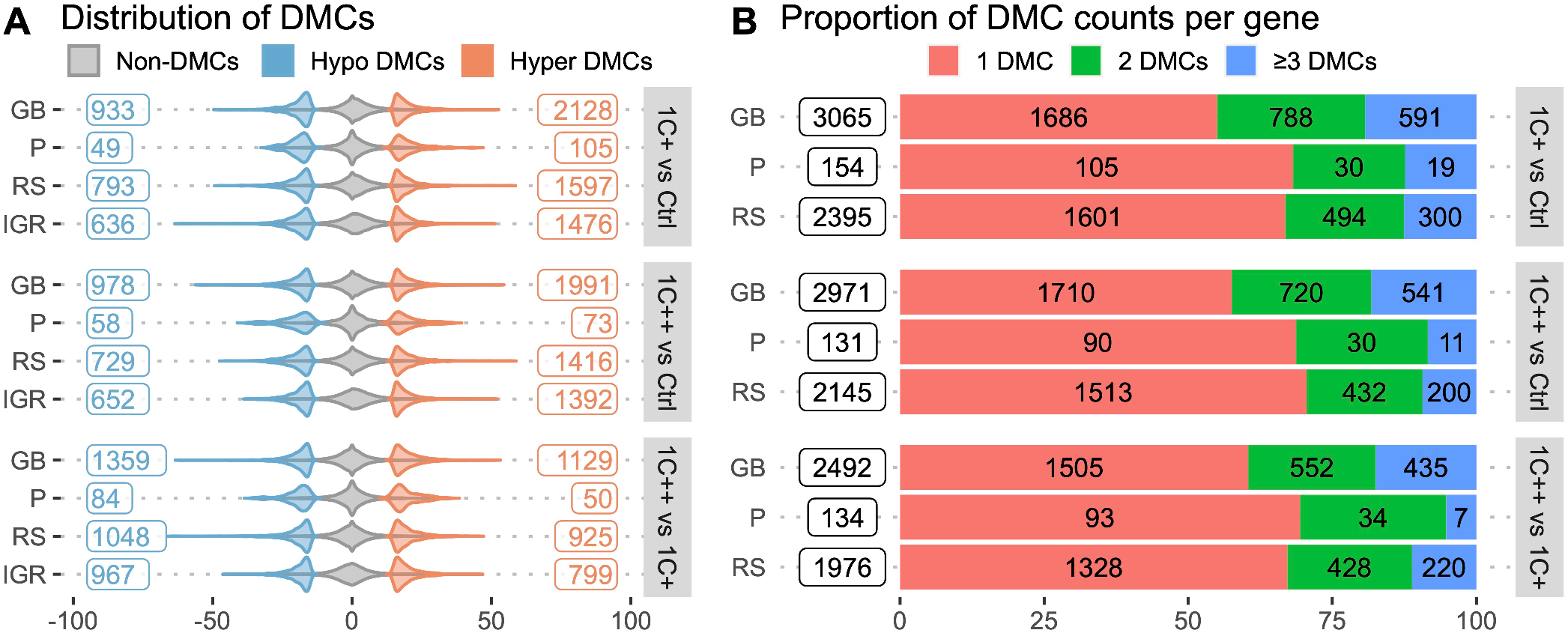
Distributions and counts of DMCs identified by three pair-wise comparisons. **(A)** Violin plots showing distributions and counts of non-DMCs, hypo DMCs, and hyper DMRs in four different genomic regions for three comparisons. The x-axis represents the percentage differences of methylation rates. **(B)** Stacked barplots showing the number of DMCs per gene, 1 DMC (red), 2 DMCs (green), and greater than or equal to 3 DMCs (blue), in three different genomic regions for three comparisons. The x-axis represents the proportions of DMC counts in percentage.

### Filtering multiple DMCs identified key genes affected by 1C supplementation

Our analysis revealed that most genes containing DMCs had only a single DMC, with more than 55%, 68%, and 66% of genes supported by single DMCs in gene bodies, promoters, and regulatory sequences, respectively (Fig 4B). While a single DMC may be sufficient to regulate gene expression, it could also be a false positive or a single nucleotide variant. Therefore, we filtered out genes that contained multiple DMCs, based on several criteria, and selected the top three genes from six different categories (see Tables S12-S14 in S1 Info, and Materials and Methods for details).

Our filtration resulted in 22 genes containing multiple DMCs in their genetic regions: promoters (P), 1K-5K upstream from the transcription start site (RS (5K)), and exons (GB) (Table 3). All genes except one (*tbx15*) contained either hypo-methylated or hyper-methylated DMCs but not both. More than a quarter of the genes (6 out of 22) were identified by both the 1C+ vs Ctrl and 1C++ vs Ctrl comparisons, indicating that 1C supplementation altered the DNA methylation of these six genes (*rn182*, *smu1-like*, *brf1*, *rnf186*, *c3*, and one uncharacterised gene) similarly regardless of the dosage levels (Table 3).

**Table 3.**
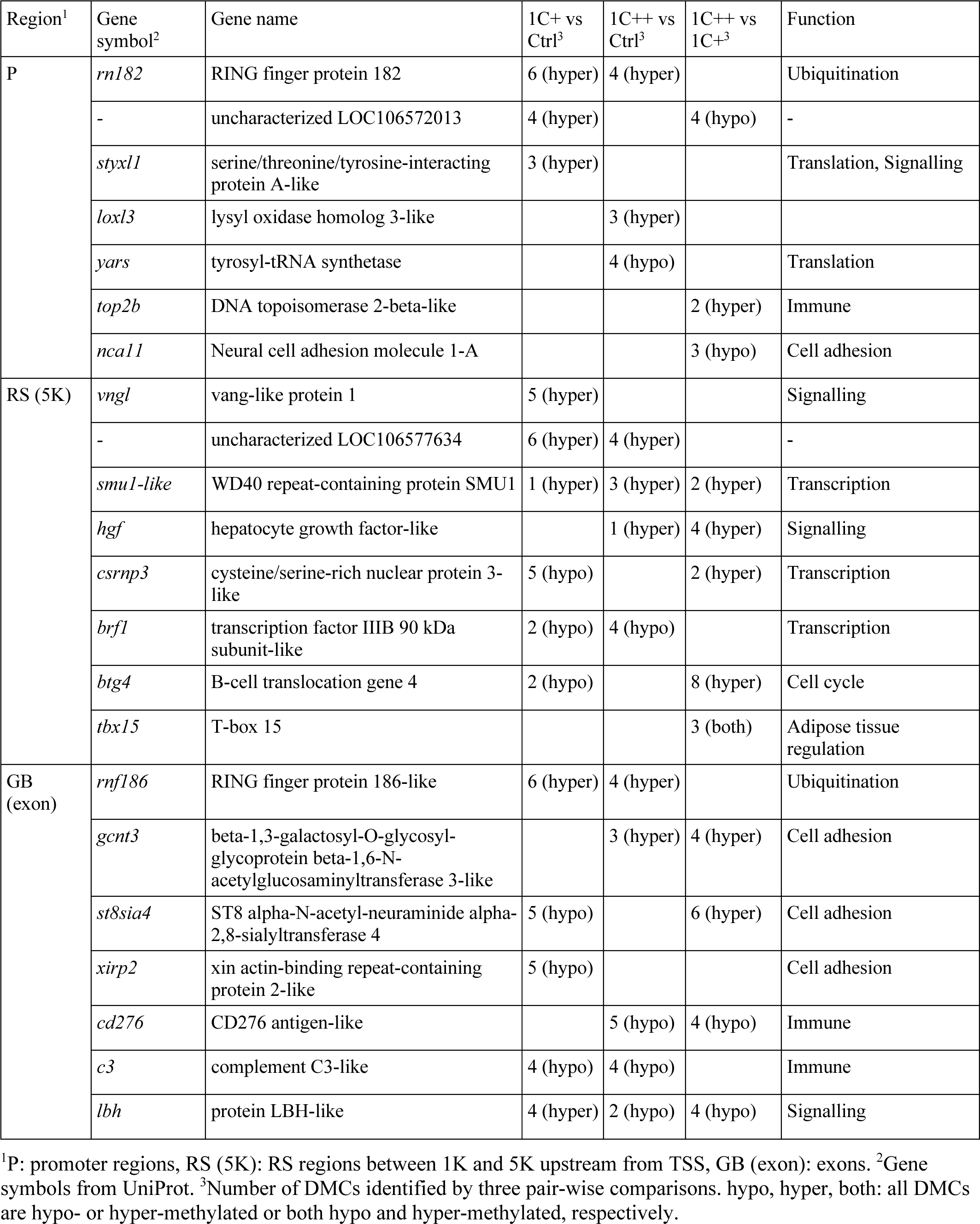
List of top three DMCs identified by three different comparisons.

Literature analysis on these 22 genes revealed that they were associated with eight biological functions: ubiquitination (*rn182* and *rnf186*) [25], translation (*styxl1* and *yars*) [26, 27], transcription (*smu1-like*, *csrnp3*, and *brf1*) [28-30], signalling (*styxl1*, *vngl*, *hgf*, and *lbh*) [26, 30], immune system (*top2b*, *cd276*, and *c3*) [31-33], cell cycle (*btg4*) [34], cell adhesion (*nca11*, *gcnt3*, *st8sia4*, and *xirp2*) [35-37], and adipose tissue regulation (*tbx15*) [38].

It is important to note that none of these 22 genes appeared to be differentially expressed at the S4 sampling point (post-smolt stage). These results suggest that 1C nutrients affected DNA methylation profiles of genes involved in various cellular processes without directly changing their gene expression levels in the liver.

### Integration of omics data revealed distinct DNA methylation landscapes in promoters and gene bodies depending on gene expression levels

To investigate the overall relationship between DNA methylation and gene expression, we integrated RNA-seq and RRBS data using all available genes and CpG sites. Most genes showed weak expression levels in the liver, as indicated by the right-skewed distribution of gene expression (Fig 5A). By considering this skewness, we grouped genes into five sets based on their normalised read counts (Fig 5B) by using four threshold values (shown as red dotted lines in Fig 5A). We also calculated average methylation rates from all 27 RRBS samples and defined six genetic regions: two each from promoters (P), regulatory sequences (RS), and gene bodies (GB).

**Fig 5.**
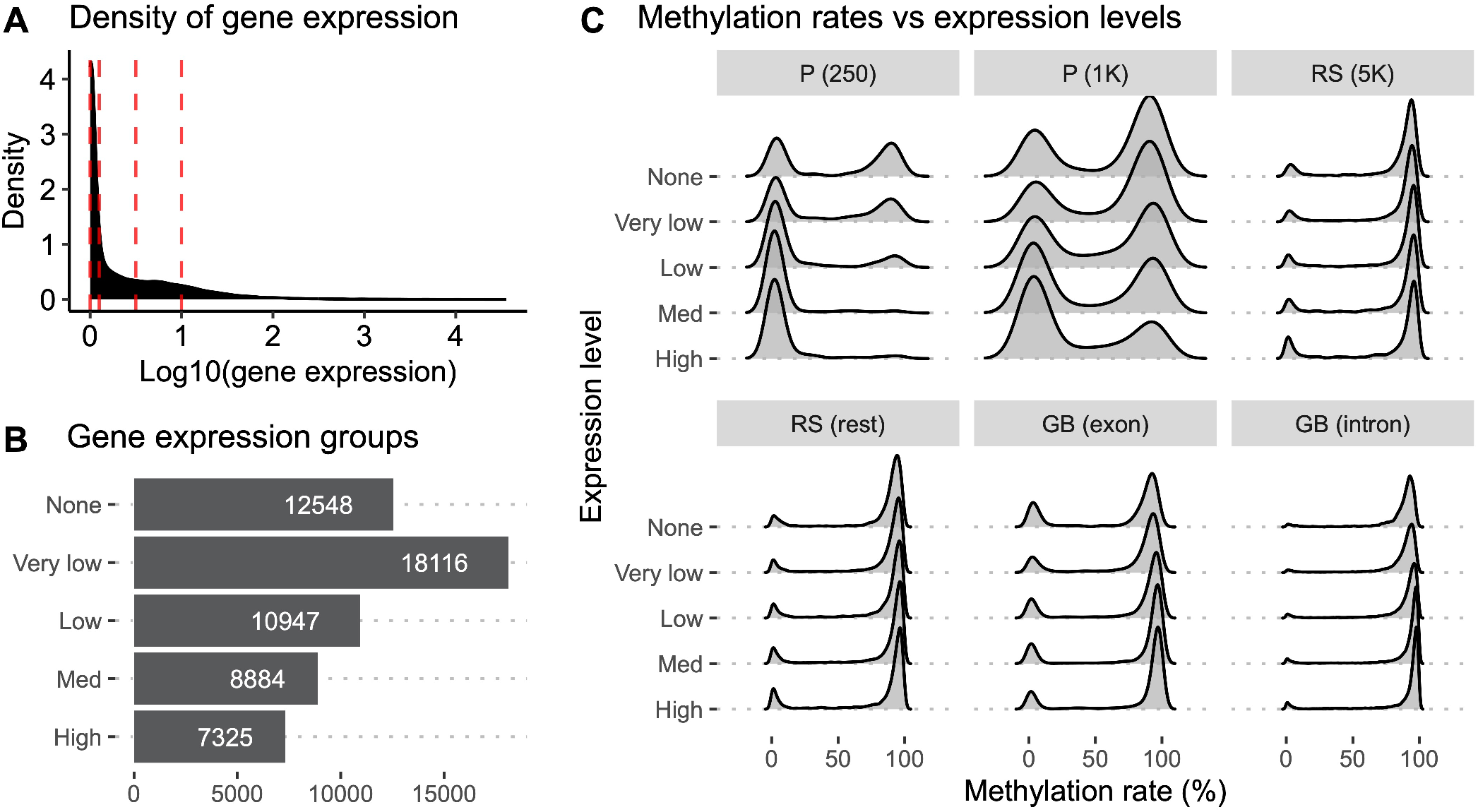
DNA methylation landscapes in five gene expression groups and six different genomic regions. **(A)** A density plot showing the distribution of gene expression calculated from normalised read counts from RNA-seq samples. Red vertical lines represent threshold values to define five different gene groups. **(B)** A barplot showing the count of genes in the five gene expression groups - None, Very low, Low, Med, and High - defined by different gene expression levels. The “None” group contains genes without any expression. **(C)** Ridge density plots displaying distributions of methylation rates for five different gene groups in six genetic regions.

Our results show that promoters and gene bodies had distinct DNA methylation landscapes depending on the expression levels of associated genes (Fig 5C). Promoter regions of higher expression genes showed lower methylation rates (0-25%), especially when located near the transcription start site (TSS). Conversely, promoter regions of genes without expression exhibited both high and low methylation rates. This trend was more pronounced when the regions were located closer to the TSS (as shown by comparing P (250) and P (1K) in Fig 5C). Furthermore, we observed that highly methylated regions (75-100%) were even more methylated (as peaks are shifted towards 100% in Fig 5C) in the bodies of genes with stronger expression levels, with a more pronounced trend observed in introns than in exons.

Overall, our findings suggest that DNA methylation landscapes vary depending on the location of the genome and the expression levels of associated genes, particularly in promoters and gene bodies. These results provide insight into the complex interplay between DNA methylation and the regulation of gene expression regulation.

### Both hypo- and hyper-methylated regions showed complex relationships with gene expression in promoters and exons

To specifically identify genes with significant differences in gene expression and DNA methylation induced by 1C nutrients, we calculated differentially methylated regions (DMRs) and selected those located within 1000 up and downstream from TSSs. This approach identified nine genes that were differentially expressed with differentially methylated regions around TSS (Table 4, Table S15 in in S1 Info).

**Table 4.**
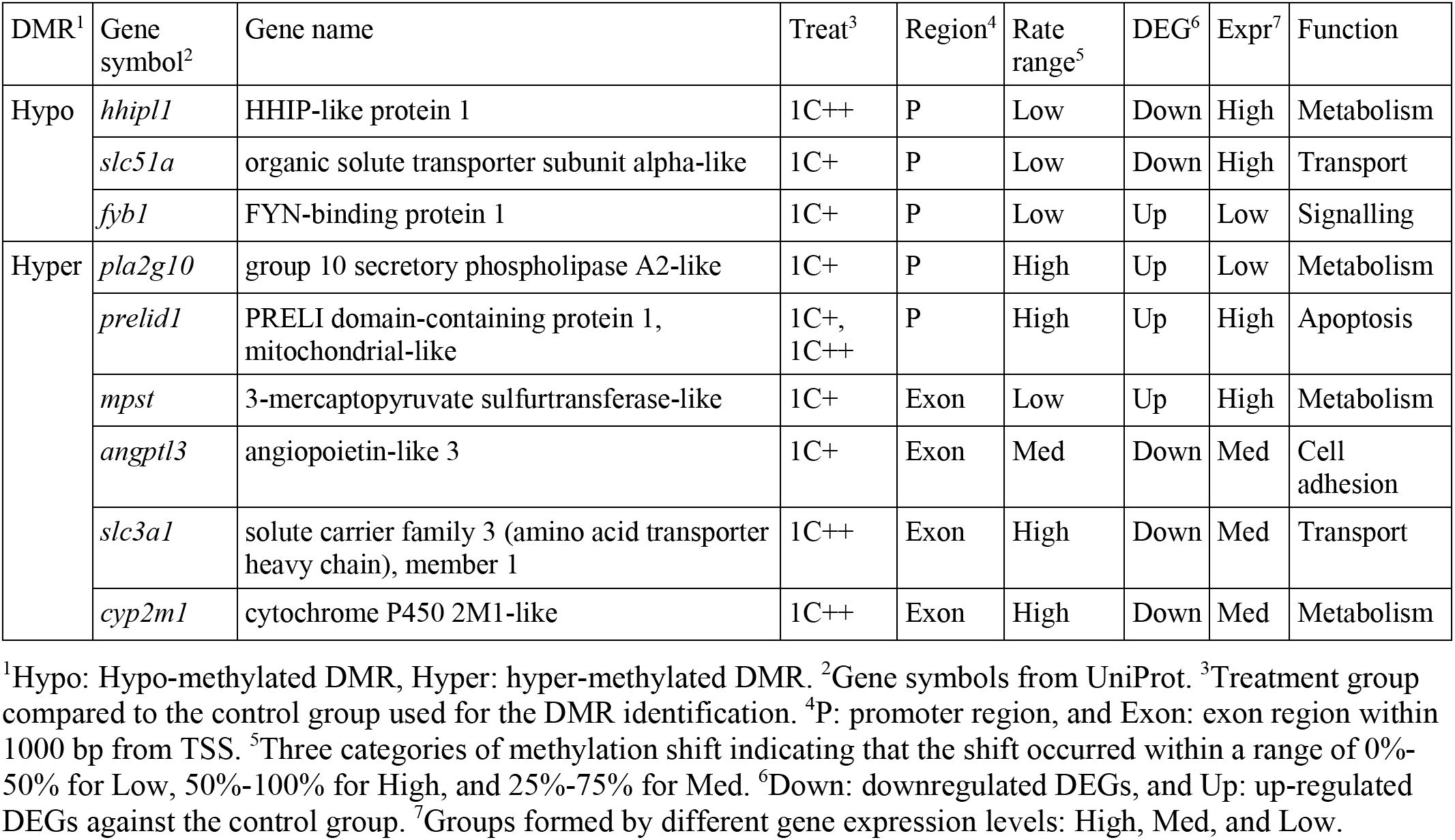
List of genes that have DMRs around their TSSs.

These genes were both hypo- and hyper-methylated in their promoters, but only hyper-methylated in the exons. In promoters, the methylation shifts occurred in a lower range of methylation rates (<50% or shown as “Low” in the Rate range column in Table 4) for the hypo-methylated DMRs, while they occurred in a higher range (>50%) for the hyper-methylated DMRs. However, in exons, the shifts of methylation occurred in a wide range of methylation rates. In addition, gene expression levels varied among the nine genes (the Expr column in Table 4). Notably, these nine genes displayed no common patterns in terms of positive or negative correlation between gene expression and DNA methylation in the liver.

In addition, literature analysis on the nine genes revealed that they were associated with five biological functions: metabolism (*hhipl1*, *pla2g10*, *mpst*, and *cyp2m1*) [30, 39, 40], cell adhesion (*angptl3*) [41], signalling (*fyb1*) [42], transport (*slc51a* and *slc3a1*) [43, 44], and apoptosis (*prelid1*) [45].

These DMRs were potentially involved in several different categories of epigenetic regulations, suggesting that they may have a complex role in gene expression rather than simply functioning as canonical on-off switches of DNA methylation in animals.

### Our analysis of omics data has generated valuable resources for future studies in nutritional epigenetics

To ensure easy access to these resources, we have prepared seven data files in a tabular format, which have been uploaded to a public data repository (https://figshare.com/s/9ab1ed68cfa1debe29d4). Additionally, basic descriptions of the data and columns can be found in Tables S16-S22 in S1 Info.

## Discussion

The growth performance of Atlantic salmon was highest with the medium dosage of 1C nutrients (1C+), followed by the high dosage (1C++) and the control diet [2]. Notably, both 1C supplementation induced significant changes in gene expression across a wide range of biological pathways in a similar manner in the liver. Specifically, the supplements affected pathways related to metabolism, genetic information processing, and cellular processes. However, up- and down-regulation of the affected genes varied within related pathways. For instance, while both supplements led to down-regulation of genes involved in steroid biosynthesis, they induced up-regulation of genes related to linoleic acid metabolism, even though both pathways are part of the lipid metabolism. Similarly, inconsistencies were found in pathways related to cytochrome P450 and translation. Moreover, the 1C supplements had an interesting effect on the amino-acid biosynthesis pathways and protein synthesis. Specifically, the supplements appeared to reduce the biosynthesis of certain amino acids while increasing protein synthesis. This suggests that the liver tissues could enhance protein synthesis without needing to activate specific amino-acid biosynthetic pathways when 1C supplements were provided.

Although most genes significantly affected by the 1C supplements showed similar expression patterns for both 1C-supplemented groups, certain genes, such as *mat2*, *lnx*, and *cyp27b1*, exhibited divergent patterns. The 1C supplements reduced the expression of two *mat2* paralogs, which were more down-regulated in the 1C++ group than the 1C+ group. Since *mat2* encodes a rate-limiting enzyme for SAM synthesis [14], the down-regulation of *mat2* could be associated with the low methylation capacity of the 1C++ group, as indicated by the SAM/SAH ratio. The 1C supplements also reduced the expression of two *lnx* paralogs, with the 1C+ group being more down-regulated than the 1C++ group. The *lnx* gene encodes an E3 ubiquitin ligase, which plays a key role in transferring ubiquitin to various protein substrates [22]. Ubiquitination is a tightly regulated post-translational modification that marks proteins for a wide range of functions, including protein degradation and subcellular localization. Furthermore, n-terminal methionine-linked ubiquitin, or linear ubiquitin, is a significant post-translational modification involved in innate immune signalling [46]. Lastly, the *cyp27b1* gene, which encodes a cytochrome P450 enzyme, exhibited the highest expression level in the 1C++ group, followed by the control and 1C+ groups. This enzyme plays an important role in calcium homeostasis by catalysing the active form of vitamin D [23]. Moreover, many other enzymes belonging to the P450 superfamily are related to cholesterol and steroid regulations [47]. In summary, these three genes and their associated regulations, such as the methylation capacity, ubiquitination, and calcium homeostasis, are potential contributors to the better growth observed in the 1C+ group compared to the 1C++ group.

While most DNA methylation markers are established during early developmental stages and become stable afterward, some markers are dynamic throughout the entire life stage [7, 48]. Our DNA methylation analysis revealed that both hypo- and hyper-methylation markers appeared to be well-conserved, as we observed very few methylation markers shifting from low methylation (0-25%) to high methylation (75%-100%) or vice versa. However, we also noticed that already methylated sites became even more methylated with 1C supplementation in the same order as the observed growth performance. This suggests that these shifted markers can be functional as DNA methylation markers for assessing growth performance.

Gene annotation of DMCs and filtering with DMC counts led us to identify 22 genes with multiple DMCs. Functional annotation of these genes revealed that they were associated with seven different biological categories: ubiquitination, translation, transcription, signalling, immune system, cell adhesion, and adipose tissue regulation. Among these categories, we focused on ubiquitination and translation, as their gene expression levels were also affected by the 1C supplements. For genes associated with ubiquitination, we observed hyper-methylation of two RING finger genes (*rn182* and *rnf186*) and significant down-regulation of gene expression in two other genes (*lnx* and *lnx1*). RING finger genes and *lnx* homologues are all E3 ubiquitin ligases linked to ubiquitination [22, 25]. For genes associated with translation, we identified two enzyme-encoding genes (*styxl1* and *yars*). The *styxl1* gene plays crucial roles in several cellular functions, including protein synthesis, signal transduction, and cell growth [26], while the *yars* gene encodes tyrosyl-tRNA synthetase that catalyses the aminoacylation of tRNA [27]. Notably, our KEGG analysis revealed that the expression levels of genes involved in the aminoacylation of tRNA were significantly down-regulated in the 1C++ group, and the promoter of the *yars* gene was hypo-methylated in the same group. These findings suggest that the 1C supplements influenced both gene expression and DNA methylation status of the genes involved in protein synthesis and ubiquitination.

Instead of limiting our analysis to genes showing significant differences, such as DEGs and genes with DMCs or DMRs, we took a more comprehensive approach by studying DNA methylation across various genomic regions and gene expression levels, by utilising the whole datasets. Our findings highlighted significant differences in methylation patterns, particularly in promoters, and some variations in gene bodies. Specifically, we observed that genes with higher expression levels exhibit lower methylation rates near their transcription start sites, while their gene bodies tend to have increased methylation in regions that were already highly methylated. These findings have raised interesting questions about the relationship between DNA methylation and gene expression. While it is commonly believed that methylation in promoters is linked to gene repression and that hypo-methylation in promoters and hyper-methylation in gene bodies are associated with gene expression [7], these correlations may not always be valid across different regions and gene expression levels. It is important to note that this hypothesis may have a strong impact on how the results of nutritional experiments should be interpreted as differences among treatment groups of such experiments can be subtle. Therefore, it is crucial to provide additional information when combining gene expression and DNA methylation data instead of solely relying on correlation coefficients for filtering.

Prior to combining two omics data sets, we implemented two pre-processing steps: identification of DMRs and filtering of DMRs to only include those around TSSs. This reduced noise and restricted the number of candidate genes whose gene expression and DNA methylation were significantly influenced by 1C supplements. By merging the two omics data sets, we identified nine genes associated with various biological functions, such as metabolism, signalling, cell adhesion, and transport. We did not observe any consistent correlation patterns between gene expression and DNA methylation for these genes, suggesting their potential involvement in complex transcriptional regulation through enhancers, silencers, and chromatin remodelling. Furthermore, some of these genes were implicated in multiple biological pathways. For example, *pla2g10* and *mpst*, two enzymes linked to metabolism, were associated with a wide range of KEGG pathways, including cysteine and methionine metabolism, and linoleic acid metabolism [30]. The *slc51a* gene is responsible for bile acid transport but also plays a role in steroid transport [43], while the *cyp2m1* gene encodes a cytochrome P450 enzyme with fatty acid hydroxylation activity [39].

Our study shares noticeable similarities with a previous salmon study that investigated the effects of varying levels of 24 micronutrients on growth performance [49]. Despite differences in feeding trials, both studies found a positive association between the down-regulation of steroid synthesis in the liver and improved growth. Additionally, we observed dynamic methylation alterations in CpG sites associated with genes involved in cell signalling and cell adhesion, which is consistent with the previous study’s findings. Furthermore, our results align with another study that examined metabolites and gene expression in skeletal muscle tissues from the same feeding trial of 1C nutrients [50]. This study demonstrated that the down-regulation of amino acid biosynthesis was associated with reduced levels of free amino acids in the muscle tissues of the 1C-supplemented group.

The effects of 1C nutrients go beyond their role in fatty acid metabolism and cell adhesion and signalling. Specifically, we found that the groups supplemented with 1C nutrients had the ability to bypass the need for synthesizing certain essential amino acids, such as methionine, and allocate more resources towards protein synthesis. As unnecessary and damaged proteins are tagged by ubiquitin for degradation [51], the control group potentially required recycling of proteins more than the 1C-supplemented groups. Additionally, a high dosage (1C++) had a negative impact on methylation capacity. Although the underlying mechanism is unclear, the difference of DNA methylation patterns between two 1C-supplemented groups may have been associated with better growth performance of the medium dosage group. Furthermore, only a few genes showed significant differences in expression and DNA methylation at the same time, many affected genes were associated with the same underlying biological pathways and regulations.

Our findings provide valuable insights into the complex regulation of gene expression and DNA methylation that are associated with growth performance, particularly with regard to protein synthesis. These insights could have significant implications for developing strategies to enhance growth and improve health outcomes from both genetic and epigenetic perspectives in salmon aquaculture, as well as in a wide range of nutrient studies.

## Materials and Methods

### Feeding trial and sampling

The details of experimental design and sampling methods were described elsewhere [2]. In brief, triplicate groups of Atlantic salmon were fed three diets for six weeks in fresh water, through smoltification, followed by three months on-growing period in salt water. At the end of the saltwater period, liver samples were dissected for RNA and DNA extraction. The samples were immediately frozen in liquid nitrogen. Fish were anaesthetised with Tricaine (Pharmaq) before any handling. All experimental procedures and husbandry practices were conducted in compliance with the Norwegian Regulation on Animal Experimentation and European Community Directive 86/609/EEC.

### Growth measurement

All fish per tank were individually measured for body weight and fork length. Following measurement, all fish were allowed to recover in aerated water. Fulton’s condition factor (K) was calculated using K = (W / L^3^) * 100 where W is body weight (g), and L is fork length (mm). Hepatosomatic index (HSI) was calculated as HSI (%) = (liver weight (g) / body weight (g)) * 100. More detailed methods along with the results of growth performance have been presented elsewhere [2].

### SAM/SAH analysis

SAM and SAH were analysed on reverse HPLC following deproteinization in 0.4M HClO4 as described by Wang et al. [52]. Five samples from each tank were pooled and homogenised for both SAM and SAH analysis.

### DNA and RNA extraction

RNA was extracted using the BioRobot EZ1 and EZ1 RNA Universal Tissue kit (Qiagen) and DNase-treated with Ambion DNA-free DNA removal kit (Invitrogen, USA) according to the manufacturer’s protocols. RNA quantity and quality were assessed using NanoDrop ND-1000 Spectrophotometer (Nanodrop Technologies) and Agilent 2100 Bioanalyser (RNA 6000 Nano LabChip kit, Agilent Technologies).

DNA isolation was performed using the DNeasy Blood & Tissue Kit (Qiagen, Cat. No. #69506) according to the manufacturer’s protocol. Liver samples were pre-treated with RNase A (provided in the Qiagen kit, 50ng/µL, 10 min at room temperature) immediately followed by proteinase K treatment (New England biolabs, #8102S 20µg/µL, 1.5 h at 55°C). DNA was eluted in Milli Q water, and its quantification was performed using the Qubit High Sensitivity Assay (Life Technologies #Q32854).

### Atlantic salmon genome and genomic annotation

We downloaded the reference genome (ICSASG version 2) and RefSeq data (version 100) for gene annotation and coordinates from the NCBI site (https://www.ncbi.nlm.nih.gov/assembly/GCF_000233375.1). All genome regions were annotated by four genetic regions as regulatory sequence (RS), promoter (P), gene body (GB), and intergenic region (IGR). Gene symbols from UniProt [53] were used when gene symbols were not available at NCBI.

### Library preparation of high-throughput sequencing

Liver samples were sent to the DeepSeq sequencing facility at Nord University, Bodø, Norway for RNA-seq. The library preparation was done using an NEBNext Ultra II Directional RNA Library Prep Kit for Illumina (New England Biolabs). The libraries were sequenced on the NextSeq500 instrument (Illumina). Detailed methods were described elsewhere [49].

Liver samples were sent to the CeMM Biomedical Sequencing Facility, Vienna, Austria for RRBS. Extracted genomic DNA were digested by MspI prior to size selection and bisulfite conversion. RRBS libraries were sequenced on Illumina HiSeq 3000/4000 instruments. Detailed methods were described elsewhere [49].

### Pre-processing of high-throughput sequencing

We used the same procedures of pre-processing for RNA-seq and RRBS data as described in a previous study [49]. In short, reads were trimmed by Cutadapt [54] and Trim Galore! (Barbraham Institute). Trimmed reads were aligned to the reference genome by STAR [55] with the default parameters for RNA-seq and by Bismark [56] and Bowtie 1 [57] with the default parameters for RRBS. While the mapped reads of RNA-seq were quantified by featureCounts [58], the mapped reads of RRBS were processed by Bismark for methylation calling and CpG site extraction. A total of 27 samples were divided into three diet groups (n=9 per group) named, 1C+, 1C++, and Ctrl, defined by different 1C nutrient levels (Fig 1 & Table S1 in S1 Info) for both RNA-seq and RRBS. The factoextra package (https://CRAN.R-project.org/package=factoextra) was used to perform clustering analysis including PCA prior to further analysis. A variance stabilizing transformation (VST) was performed on RNA-seq counts using DESeq2 [59] prior to PCA. A non-linear clustering method, t-distributed stochastic neighbor embedding (t-SNE) [60], was additionally used for RRBS.

### Differential gene expression analysis

Differentially expressed genes (DEGs) of three comparisons (1C+ vs Ctrl, 1C++ vs Ctrl, and 1C++ vs 1C+) were identified by DESeq2 [59] when the adjusted p-values by Benjamini-Hochberg [61] were less than 0.05. To find DEGs strongly affected by the treatment, we used log fold changes (LFCs) from pair-wise comparisons for filtering by two thresholds, |LFC| > 1.5 and |LFC| > 2. The LFC filtration was performed by using the lfcThreshold argument of the results function of DESeq2 [59].

For the clustering analysis with density-based spatial clustering of applications with noise (DBSCAN) [62], the results of the three comparisons were combined into a pooled set of DEGs without redundancy. The input of DBSCAN were pair-wise differences of group-wise gene expression levels calculated from the pooled set of DEGs. To calculate the average gene expression levels for each group, the trimmed mean of M-values (TMM) normalization was calculated by edgeR [63] and then scaled by the minimum and maximum values to the range between 0 and 1. The DBSCAN clustering was performed by the dbscan package (https://CRAN.R-project.org/package=dbscan) with parameters minPts=100 and eps=0.5.

### Functional annotation with KEGG

Both over-representation analysis (ORA) and gene set enrichment analysis (GSEA) [64] on the Kyoto Encyclopedia of Genes and Genomes (KEGG) database [30] were performed by clusterProfiler [65]. The input of ORA were gene lists of DBSCAN clusters, whereas that of GSEA was shrunken LFCs calculated by the normal shrinkage method provided by DESeq2 [59], being separately calculated for three pair-wise comparisons. P-values were adjusted by Benjamini-Hochberg [61]. Enriched pathways were identified when adjusted p-values were less than 0.05 for ORA and 0.01 for GSEA. For GSEA, the enriched pathways were selected when supported by at least two results.

### Differential methylation analysis

Prior to differential methylation analysis, reads with less than or equal to 10 and above the 99.9^th^ percentile of coverage were discarded by methylKit [66]. Differentially methylated CpG sites (DMCs) of three comparisons (1C+ vs Ctrl, 1C++ vs Ctrl, and 1C++ vs 1C+) were identified by methylKit [66] when the Q-values by the sliding linear model (SLIM) method [67] were less than 0.05 and methylation differences greater than or equal to 15%.

To filter out DMCs that were strongly affected by the treatments, we used the counts of DMCs per gene in three different regions: promoter, regulatory regions within 5000 upstream from transcription start sites (TSSs), and exons. Genes were sorted by the count of DMCs in a descendant order, and the top three genes among them were selected for the further literature analysis. Genes with higher average methylation differences were selected in the case of ties.

To identify differentially methylated regions (DMRs), average methylation rates were calculated by sliding windows with size 100 and step 100 using the tileMethylCounts function of methylKit [66]. The same procedure of identifying DMCs was applied to identify DMRs after tiled regions were defined.

### Linking DNA methylation with gene expression

To investigate a broad association between gene expression and DNA methylation, we divided genes into five gene groups - None, Very low, Low, Med, and High, by gene expression levels, which were defined from scaled TMM counts. Threshold ranges used to define the groups were 0 for None, (0, 0.1], (0.1, 0.5], and (0.5, 1] for Very low, Low, Med, respectively, and >1 for High. Linking DEGs with DMCs or DMRs was done by merging NCBI gene IDs from both datasets.

### Bioinformatics analysis

In-house R and Python scripts with Snakemake [68] were used to perform high-throughput sequence analysis, basic statistical analysis, and generating figures.

## Acknowledgements

We thank Amelie Nemc and Bekir Ergüner for advice on RRBS analysis, and Hui-Shan Tung and Eva Mykkeltvedt for DNA and RNA extraction at the Institute of Marine Research (Bergen, Norway). This work was supported by the Norwegian Research Council under project no: 267787 (NutrEpi) and by the Institute of Marine Research under the Nutritional programming project.

## Supporting Information

S1 Info contains 22 supplementary tables and one supplementary figure.

**Table S1**. Composition of the experimental diets (g/kg).

**Table S2**. Growth performance measured at four sampling points.

**Table S3**. Read counts of RNA-seq samples after initial quality control, alignment, and quantification.

**Table S4**. Number of DEGs identified by three comparisons.

**Table S5**. Enriched KEGG pathways for the genes in DEG C1 and DEG C2 clusters by ORA.

**Table S6**. Enriched KEGG pathways for the C1+ vs Ctrl comparison by GSEA.

**Table S7**. Enriched KEGG pathways for the C1++ vs Ctrl comparison by GSEA.

**Table S8**. Enriched KEGG pathways for the C1+ vs Ctrl comparison by GSEA.

**Table S9**. Read counts of RRBS samples after initial quality control and alignment percentage.

**Table S10**. Comparisons of two methylation rate distributions in different regions by KS tests.

**Table S11**. Number of DMCs identified by three comparisons in four different regions.

**Table S12**. List of genes that have multiple DMCs in the promoter (P) regions.

**Table S13**. List of genes that have multiple DMCs in the RS (5K) regions.

**Table S14**. List of genes that have multiple DMCs in the GB (exon) regions.

**Table S15**. List of genes that are DEGs and contain DMRs around their TSSs.

**Table S16**. List of DEGs from three comparisons: 1C+ vs Ctrl, 1C++ vs Ctrl, and 1C++ vs 1C+.

**Table S17**. List of DEGs from three comparisons: 1C+ vs Ctrl, 1C++ vs Ctrl, and 1C++ vs 1C+.

**Table S18**. List of DEGs from three comparisons: 1C+ vs Ctrl, 1C++ vs Ctrl, and 1C++ vs 1C+.

**Table S19**. List of DEG clusters identified by DBSCAN.

**Table S20**. List of enriched KEGG pathways identified by ORA and GSEA.

**Table S21**. List of DMCs from three comparisons: 1C+ vs Ctrl, 1C++ vs Ctrl, and 1C++ vs 1C+.

**Table S22**. List of DMRs from three comparisons: 1C+ vs Ctrl, 1C++ vs Ctrl, and 1C++ vs 1C+.

**Fig S1**. Clustering analysis on the methylation rates of mapped CpG.

## References

1. Ducker GS, Rabinowitz JD. One-Carbon Metabolism in Health and Disease. Cell Metab. 2017;25(1):27–42. doi: 10.1016/j.cmet.2016.08.009. PubMed PMID: WOS:000392845500007.

2. Espe M, Vikesa V, Thomsen TH, Adam AC, Saito T, Skjaerven KH. Atlantic salmon fed a nutrient package of surplus methionine, vitamin B12, folic acid and vitamin B6 improved growth and reduced the relative liver size, but when in excess growth reduced. Aquacult Nutr. 2020;26(2):477–89. doi: 10.1111/anu.13010. PubMed PMID: WOS:000494524000001.

3. Lyon P, Strippoli V, Fang B, Cimmino L. B Vitamins and One-Carbon Metabolism: Implications in Human Health and Disease. Nutrients. 2020;12(9). doi: ARTN 2867 10.3390/nu12092867. PubMed PMID: WOS:000580091900001.

4. Danchin A, Sekowska A, You CH. One-carbon metabolism, folate, zinc and translation. Microb Biotechnol. 2020;13(4):899–925. doi: 10.1111/1751-7915.13550. PubMed PMID: WOS:000527739000001.

5. Hamre K, Sissener NH, Lock EJ, Olsvik PA, Espe M, Torstensen BE, et al. Antioxidant nutrition in Atlantic salmon (Salmo salar) parr and post-smolt, fed diets with high inclusion of plant ingredients and graded levels of micronutrients and selected amino acids. Peerj. 2016;4. doi: ARTN e2688 10.7717/peerj.2688. PubMed PMID: WOS:000387171900012.

6. Hemre GI, Lock EJ, Olsvik PA, Hamre K, Espe M, Torstensen BE, et al. Atlantic salmon (Salmo salar) require increased dietary levels of B-vitamins when fed diets with high inclusion of plant based ingredients. Peerj. 2016;4. doi: ARTN e2493 10.7717/peerj.2493. PubMed PMID: WOS:000385572500001.

7. Lanata CM, Chung SA, Criswell LA. DNA methylation 101: what is important to know about DNA methylation and its role in SLE risk and disease heterogeneity. Lupus Sci Med. 2018;5(1). doi: UNSP e000285 10.1136/lupus-2018-000285. PubMed PMID: WOS:000495994400032.

8. Xu J, Sinclair KD. One-carbon metabolism and epigenetic regulation of embryo development. Reprod Fert Develop. 2015;27(4):667–76. doi: 10.1071/Rd14377. PubMed PMID: WOS:000353147100010.

9. Brosnan JT, Jacobs RL, Stead LM, Brosnan ME. Methylation demand: a key determinant of homocysteine metabolism. Acta Biochim Pol. 2004;51(2):405–13. PubMed PMID: WOS:000222272700012.

10. Arslan C. L-carnitine and its use as a feed additive in poultry feeding a review. Rev Med Vet-Toulouse. 2006;157(3):134–42. PubMed PMID: WOS:000237402400003.

11. Noga AA, Vance DE. Insights into the requirement of phosphatidylcholine synthesis for liver function in mice. J Lipid Res. 2003;44(10):1998–2005. doi: 10.1194/jlr.M300226-JLR200. PubMed PMID: WOS:000186043900021.

12. da Silva RP, Kelly KB, Al Rajabi A, Jacobs RL. Novel insights on interactions between folate and lipid metabolism. Biofactors. 2014;40(3):277–83. doi: 10.1002/biof.1154. PubMed PMID: WOS:000337634300001.

13. Casero RA, Pegg AE. Polyamine catabolism and disease. Biochem J. 2009;421:323-38. doi: 10.1042/Bj20090598. PubMed PMID: WOS:000268615900001.

14. Stipanuk MH. Metabolism of Sulfur-Containing Amino Acids: How the Body Copes with Excess Methionine, Cysteine, and Sulfide. J Nutr. 2020;150:2494s–505s. doi: 10.1093/jn/nxaa094. PubMed PMID: WOS:000579421800002.

15. Finkelstein JD, Martin JJ. Homocysteine. Int J Biochem Cell Biol. 2000;32(4):385–9. Epub 2000/04/13. doi: 10.1016/s1357-2725(99)00138-7. PubMed PMID: 10762063.

16. Finkelstein JD. Methionine Metabolism in Mammals. J Nutr Biochem. 1990;1(5):228–37. doi: Doi 10.1016/0955-2863(90)90070-2. PubMed PMID: WOS:A1990DD46200001.

17. Torstensen BE, Espe M, Stubhaug I, Lie O. Dietary plant proteins and vegetable oil blends increase adiposity and plasma lipids in Atlantic salmon (Salmo salar L.). Brit J Nutr. 2011;106(5):633–47. doi: 10.1017/S0007114511000729. PubMed PMID: WOS:000294882000002.

18. Espe M, Rathore RM, Du ZY, Liaset B, El-Mowafi A. Methionine limitation results in increased hepatic FAS activity, higher liver 18:1 to 18:0 fatty acid ratio and hepatic TAG accumulation in Atlantic salmon, Salmo salar. Amino Acids. 2010;39(2):449–60. doi: 10.1007/s00726-009-0461-2. PubMed PMID: WOS:000279190400015.

19. Vanni E, Bugianesi E, Kotronen A, De Minicis S, Yki-Jarvinen H, Svegliati-Baroni G. From the metabolic syndrome to NAFLD or vice versa? Digest Liver Dis. 2010;42(5):320–30. doi: 10.1016/j.dld.2010.01.016. PubMed PMID: WOS:000278045500003.

20. Vernon G, Baranova A, Younossi ZM. Systematic review: the epidemiology and natural history of non-alcoholic fatty liver disease and non-alcoholic steatohepatitis in adults. Aliment Pharm Ther. 2011;34(3):274–85. doi: 10.1111/j.1365-2036.2011.04724.x. PubMed PMID: WOS:000292394800002.

21. Rolo AP, Teodoro JS, Palmeira CM. Role of oxidative stress in the pathogenesis of nonalcoholic steatohepatitis. Free Radical Bio Med. 2012;52(1):59–69. doi: 10.1016/j.freeradbiomed.2011.10.003. PubMed PMID: WOS:000298975500006.

22. Doyle JM, Gao JL, Wang JW, Yang MJ, Potts PR. MAGE-RING Protein Complexes Comprise a Family of E3 Ubiquitin Ligases. Mol Cell. 2010;39(6):963–74. doi: 10.1016/j.molcel.2010.08.029. PubMed PMID: WOS:000282377200015.

23. Zehnder D, Bland R, Williams MC, McNinch RW, Howie AJ, Stewart PM, et al. Extrarenal expression of 25-hydroxyvitamin d(3)-1 alpha-hydroxylase. J Clin Endocrinol Metab. 2001;86(2):888–94. Epub 2001/02/07. doi: 10.1210/jcem.86.2.7220. PubMed PMID: 11158062.

24. Kerr SJ. Competing Methyltransferase Systems. J Biol Chem. 1972;247(13):4248-&. PubMed PMID: WOS:A1972M928700022.

25. Lipkowitz S, Weissman AM. RINGs of good and evil: RING finger ubiquitin ligases at the crossroads of tumour suppression and oncogenesis. Nat Rev Cancer. 2011;11(9):629–43. Epub 2011/08/25. doi: 10.1038/nrc3120. PubMed PMID: 21863050; PubMed Central PMCID: PMCPMC3542975.

26. Ardito F, Giuliani M, Perrone D, Troiano G, Lo Muzio L. The crucial role of protein phosphorylation in cell signaling and its use as targeted therapy (Review). Int J Mol Med. 2017;40(2):271–80. Epub 2017/06/29. doi: 10.3892/ijmm.2017.3036. PubMed PMID: 28656226; PubMed Central PMCID: PMCPMC5500920.

27. Ribas de Pouplana L, Frugier M, Quinn CL, Schimmel P. Evidence that two present-day components needed for the genetic code appeared after nucleated cells separated from eubacteria. Proc Natl Acad Sci U S A. 1996;93(1):166–70. Epub 1996/01/09. doi: 10.1073/pnas.93.1.166. PubMed PMID: 8552597; PubMed Central PMCID: PMCPMC40199.

28. Keiper S, Papasaikas P, Will CL, Valcarcel J, Girard C, Luhrmann R. Smu1 and RED are required for activation of spliceosomal B complexes assembled on short introns. Nat Commun. 2019;10(1):3639. Epub 2019/08/15. doi: 10.1038/s41467-019-11293-8. PubMed PMID: 31409787; PubMed Central PMCID: PMCPMC6692369.

29. Wang Z, Roeder RG. Structure and function of a human transcription factor TFIIIB subunit that is evolutionarily conserved and contains both TFIIB- and high-mobility-group protein 2-related domains. Proc Natl Acad Sci U S A. 1995;92(15):7026–30. Epub 1995/07/18. doi: 10.1073/pnas.92.15.7026. PubMed PMID: 7624363; PubMed Central PMCID: PMCPMC41464.

30. Kanehisa M, Goto S. KEGG: Kyoto Encyclopedia of Genes and Genomes. Nucleic Acids Res. 2000;28(1):27–30. doi: DOI 10.1093/nar/28.1.27. PubMed PMID: WOS:000084896300007.

31. Papapietro O, Chandra A, Eletto D, Inglott S, Plagnol V, Curtis J, et al. Topoisomerase 2beta mutation impairs early B-cell development. Blood. 2020;135(17):1497–501. Epub 2020/03/05. doi: 10.1182/blood.2019003299. PubMed PMID: 32128574; PubMed Central PMCID: PMCPMC7612232.

32. Sahu A, Lambris JD. Structure and biology of complement protein C3, a connecting link between innate and acquired immunity. Immunol Rev. 2001;180:35–48. Epub 2001/06/21. doi: 10.1034/j.1600-065x.2001.1800103.x. PubMed PMID: 11414361.

33. Kontos F, Michelakos T, Kurokawa T, Sadagopan A, Schwab JH, Ferrone CR, et al. B7-H3: An Attractive Target for Antibody-based Immunotherapy. Clin Cancer Res. 2021;27(5):1227–35. Epub 2020/10/15. doi: 10.1158/1078-0432.CCR-20-2584. PubMed PMID: 33051306; PubMed Central PMCID: PMCPMC7925343.

34. Buanne P, Corrente G, Micheli L, Palena A, Lavia P, Spadafora C, et al. Cloning of PC3B, a novel member of the PC3/BTG/TOB family of growth inhibitory genes, highly expressed in the olfactory epithelium. Genomics. 2000;68(3):253–63. doi: 10.1006/geno.2000.6288. PubMed PMID: WOS:000089549900004.

35. Berger RP, Sun YH, Kulik M, Lee JK, Nairn AV, Moremen KW, et al. ST8SIA4-Dependent Polysialylation is Part of a Developmental Program Required for Germ Layer Formation from Human Pluripotent Stem Cells. Stem Cells. 2016;34(7):1742–52. Epub 2016/04/14. doi: 10.1002/stem.2379. PubMed PMID: 27074314; PubMed Central PMCID: PMCPMC4931981.

36. Wang Q, Lin JL, Erives AJ, Lin CI, Lin JJ. New insights into the roles of Xin repeat-containing proteins in cardiac development, function, and disease. Int Rev Cell Mol Biol. 2014;310:89–128. Epub 2014/04/15. doi: 10.1016/B978-0-12-800180-6.00003-7. PubMed PMID: 24725425; PubMed Central PMCID: PMCPMC4857591.

37. Sumardika IW, Youyi C, Kondo E, Inoue Y, Ruma IMW, Murata H, et al. beta-1,3-Galactosyl-O-Glycosyl-Glycoprotein beta-1,6-N-Acetylglucosaminyltransferase 3 Increases MCAM Stability, Which Enhances S100A8/A9-Mediated Cancer Motility. Oncol Res. 2018;26(3):431-44. Epub 2017/09/20. doi: 10.3727/096504017X15031557924123. PubMed PMID: 28923134; PubMed Central PMCID: PMCPMC7844831.

38. Pan DZ, Miao Z, Comenho C, Rajkumar S, Koka A, Lee SHT, et al. Identification of TBX15 as an adipose master trans regulator of abdominal obesity genes. Genome Med. 2021;13(1):123. Epub 2021/08/04. doi: 10.1186/s13073-021-00939-2. PubMed PMID: 34340684; PubMed Central PMCID: PMCPMC8327600.

39. Yang YH, Wang JL, Miranda CL, Buhler DR. CYP2M1: cloning, sequencing, and expression of a new cytochrome P450 from rainbow trout liver with fatty acid (omega-6)-hydroxylation activity. Arch Biochem Biophys. 1998;352(2):271–80. Epub 1998/05/20. doi: 10.1006/abbi.1998.0607. PubMed PMID: 9587416.

40. Stelzer G, Rosen N, Plaschkes I, Zimmerman S, Twik M, Fishilevich S, et al. The GeneCards Suite: From Gene Data Mining to Disease Genome Sequence Analyses. Curr Protoc Bioinformatics. 2016;54:1 30 1-1 3. Epub 2016/06/21. doi: 10.1002/cpbi.5. PubMed PMID: 27322403.

41. Camenisch G, Pisabarro MT, Sherman D, Kowalski J, Nagel M, Hass P, et al. ANGPTL3 stimulates endothelial cell adhesion and migration via integrin alpha vbeta 3 and induces blood vessel formation in vivo. J Biol Chem. 2002;277(19):17281–90. Epub 2002/03/06. doi: 10.1074/jbc.M109768200. PubMed PMID: 11877390.

42. Griffiths EK, Penninger JM. Communication between the TCR and integrins: role of the molecular adapter ADAP/Fyb/Slap. Curr Opin Immunol. 2002;14(3):317–22. Epub 2002/04/26. doi: 10.1016/s0952-7915(02)00334-5. PubMed PMID: 11973129.

43. Ballatori N, Christian WV, Lee JY, Dawson PA, Soroka CJ, Boyer JL, et al. OSTalpha-OSTbeta: a major basolateral bile acid and steroid transporter in human intestinal, renal, and biliary epithelia. Hepatology. 2005;42(6):1270–9. Epub 2005/12/01. doi: 10.1002/hep.20961. PubMed PMID: 16317684.

44. Lee WS, Wells RG, Sabbag RV, Mohandas TK, Hediger MA. Cloning and chromosomal localization of a human kidney cDNA involved in cystine, dibasic, and neutral amino acid transport. J Clin Invest. 1993;91(5):1959–63. Epub 1993/05/01. doi: 10.1172/JCI116415. PubMed PMID: 8486766; PubMed Central PMCID: PMCPMC288191.

45. Potting C, Tatsuta T, Konig T, Haag M, Wai T, Aaltonen MJ, et al. TRIAP1/PRELI complexes prevent apoptosis by mediating intramitochondrial transport of phosphatidic acid. Cell Metab. 2013;18(2):287–95. Epub 2013/08/13. doi: 10.1016/j.cmet.2013.07.008. PubMed PMID: 23931759.

46. Fiil BK, Gyrd-Hansen M. Met1-linked ubiquitination in immune signalling. Febs J. 2014;281(19):4337–50. doi: 10.1111/febs.12944. PubMed PMID: WOS:000342825100001.

47. Lorbek G, Lewinska M, Rozman D. Cytochrome P450s in the synthesis of cholesterol and bile acids - from mouse models to human diseases. Febs J. 2012;279(9):1516–33. doi: 10.1111/j.1742-4658.2011.08432.x. PubMed PMID: WOS:000302995500002.

48. Jones PA. Functions of DNA methylation: islands, start sites, gene bodies and beyond. Nat Rev Genet. 2012;13(7):484–92. doi: 10.1038/nrg3230. PubMed PMID: WOS:000305464800010.

49. Saito T, Whatmore P, Taylor JF, Fernandes JMO, Adam AC, Tocher DR, et al. Micronutrient supplementation affects transcriptional and epigenetic regulation of lipid metabolism in a dose-dependent manner. Epigenetics-Us. 2021;16(11):1217–34. doi: 10.1080/15592294.2020.1859867. PubMed PMID: WOS:000603885200001.

50. Adam AC, Saito T, Espe M, Whatmore P, Fernandes JMO, Vikesa V, et al. Metabolic and molecular signatures of improved growth in Atlantic salmon (Salmo salar) fed surplus levels of methionine, folic acid, vitamin B(6) and B(12) throughout smoltification. Br J Nutr. 2022;127(9):1289–302. Epub 2021/06/29. doi: 10.1017/S0007114521002336. PubMed PMID: 34176547.

51. Ciechanover A. Intracellular protein degradation: from a vague idea thru the lysosome and the ubiquitin-proteasome system and onto human diseases and drug targeting. Cell Death Differ. 2005;12(9):1178–90. Epub 2005/08/12. doi: 10.1038/sj.cdd.4401692. PubMed PMID: 16094394.

52. Wang W, Kramer PM, Yang SM, Pereira MA, Tao LH. Reversed-phase high-performance liquid chromatography procedure for the simultaneous determination of S-adenosyl-L-methionine and S-adenosyl-L-homocysteine in mouse liver and the effect of methionine on their concentrations. J Chromatogr B. 2001;762(1):59–65. doi: Doi 10.1016/S0378-4347(01)00341-3. PubMed PMID: WOS:000171062300008.

53. Bateman A, Martin MJ, Orchard S, Magrane M, Ahmad S, Alpi E, et al. UniProt: the Universal Protein Knowledgebase in 2023. Nucleic Acids Res. 2022. doi: 10.1093/nar/gkac1052. PubMed PMID: WOS:000893231800001.

54. Martin M. Cutadapt removes adapter sequences from high-throughput sequencing reads. 2011. 2011;17(1):3. Epub 2011-08-02. doi: 10.14806/ej.17.1.200.

55. Dobin A, Davis CA, Schlesinger F, Drenkow J, Zaleski C, Jha S, et al. STAR: ultrafast universal RNA-seq aligner. Bioinformatics. 2013;29(1):15–21. doi: 10.1093/bioinformatics/bts635. PubMed PMID: WOS:000312654600003.

56. Krueger F, Andrews SR. Bismark: a flexible aligner and methylation caller for Bisulfite-Seq applications. Bioinformatics. 2011;27(11):1571–2. doi: 10.1093/bioinformatics/btr167. PubMed PMID: WOS:000291062400018.

57. Langmead B, Trapnell C, Pop M, Salzberg SL. Ultrafast and memory-efficient alignment of short DNA sequences to the human genome. Genome Biol. 2009;10(3). doi: ARTN R25 10.1186/gb-2009-10-3-r25. PubMed PMID: WOS:000266544500005.

58. Liao Y, Smyth GK, Shi W. featureCounts: an efficient general purpose program for assigning sequence reads to genomic features. Bioinformatics. 2014;30(7):923–30. doi: 10.1093/bioinformatics/btt656. PubMed PMID: WOS:000334078300005.

59. Love MI, Huber W, Anders S. Moderated estimation of fold change and dispersion for RNA-seq data with DESeq2. Genome Biol. 2014;15(12). doi: ARTN 550 10.1186/s13059-014-0550-8. PubMed PMID: WOS:000346609500022.

60. van der Maaten L, Hinton G. Visualizing Data using t-SNE. J Mach Learn Res. 2008;9:2579–605. PubMed PMID: WOS:000262637600007.

61. Benjamini Y, Hochberg Y. Controlling the False Discovery Rate - a Practical and Powerful Approach to Multiple Testing. J R Stat Soc B. 1995;57(1):289–300. doi: DOI 10.1111/j.2517-6161.1995.tb02031.x. PubMed PMID: WOS:A1995QE45300017.

62. Ester M, Kriegel H-P, Sander J, Xu X. A Density-Based Algorithm for Discovering Clusters in Large Spatial Databases with Noise. Proc of 2nd International Conference on Knowledge Discovery and. ester1996densitybased1996. p. 226-31.

63. Robinson MD, McCarthy DJ, Smyth GK. edgeR: a Bioconductor package for differential expression analysis of digital gene expression data. Bioinformatics. 2010;26(1):139–40. doi: 10.1093/bioinformatics/btp616. PubMed PMID: WOS:000273116100025.

64. Subramanian A, Tamayo P, Mootha VK, Mukherjee S, Ebert BL, Gillette MA, et al. Gene set enrichment analysis: A knowledge-based approach for interpreting genome-wide expression profiles. P Natl Acad Sci USA. 2005;102(43):15545–50. doi: 10.1073/pnas.0506580102. PubMed PMID: WOS:000232929400051.

65. Yu GC, Wang LG, Han YY, He QY. clusterProfiler: an R Package for Comparing Biological Themes Among Gene Clusters. Omics. 2012;16(5):284–7. doi: 10.1089/omi.2011.0118. PubMed PMID: WOS:000303653300007.

66. Akalin A, Kormaksson M, Li S, Garrett-Bakelman FE, Figueroa ME, Melnick A, et al. methylKit: a comprehensive R package for the analysis of genome-wide DNA methylation profiles. Genome Biol. 2012;13(10). doi: ARTN R87 10.1186/gb-2012-13-10-R87. PubMed PMID: WOS:000313183900006.

67. Wang HQ, Tuominen LK, Tsai CJ. SLIM: a sliding linear model for estimating the proportion of true null hypotheses in datasets with dependence structures. Bioinformatics. 2011;27(2):225-31. doi: 10.1093/bioinformatics/btq650. PubMed PMID: WOS:000286215200012.

68. Koster J, Rahmann S. Snakemake-a scalable bioinformatics workflow engine. Bioinformatics. 2012;28(19):2520–2. doi: 10.1093/bioinformatics/bts480. PubMed PMID: WOS:000309687500018.

