## Supplementary material for "One-carbon metabolism nutrients impact the interplay between DNA methylation and gene expression in liver, enhancing protein synthesis in Atlantic Salmon": S1 Info

### Supplementary tables

**Table S1.** Composition of the experimental diets (g/kg).

| | Ctrl | 1C+ | 1C+ |
|---|---|---|---|
| **Wheat** | 54.22 | 50.64 | 47.05 |
| **Wheat gluten** | 132.03 | 132.03 | 132.03 |
| **Sunflower meal** | 10.0 | 10.0 | 10.0 |
| **Dehulled faba beans** | 30.0 | 30.0 | 30.0 |
| **Pea concentrate** | 150.0 | 150.0 | 150.0 |
| **Soy protein concentrate** | 240.0 | 240.0 | 240.0 |
| **Krill meal** | 20.0 | 20.0 | 20.0 |
| **Fish meal** | 120.0 | 120.0 | 120.0 |
| **Rapeseed oil** | 81.24 | 81.24 | 81.24 |
| **Fish oil** | 126.9 | 126.9 | 126.9 |
| **Water** | 11.43 | 11.84 | 12.26 |
| **DL-methionine** | **0.05** | **3.12** | **6.19** |
| **Choline** | 0.92 | 0.92 | 0.92 |
| **NRC mineral mix** | 2.0 | 2.0 | 2.0 |
| **NRC Vitamin mix** | 1.0 | 1.0 | 1.0 |
| **Vitamin B12** | **0.156** | **0.179** | **0.203** |
| **Folate** | **0.023** | **0.053** | **0.083** |
| **Vitamin B6** | **0.077** | **0.107** | **0.137** |
| **Taurine** | 2.8 | 2.8 | 2.8 |
| **Micronutrients** | 17.16 | 17.17 | 17.19 |

**Table S2.** Growth performance measured at four sampling points.

| Sampling point | Measure† | Ctrl | | 1C+ | | 1C++ | | p-value |
|---|---|---|---|---|---|---|---|---|
| **S1** | BW (g) | 31.62±0.78 | | 32.16±0.47 | | 32.74±0.35 | | 0.43 |
| **S2** | BW (g) | 85.81±3.14 | | 90.54±1.24 | | 87.59±4.64 | | 0.62 |
| | CF | 1.29±0.02 | | 1.32±0.00 | | 1.31±0.03 | | 0.51 |
| | HSI | 1.09±0.04 | | 0.99±0.00 | | 1.02±0.01 | | 0.06 |
| **S3** | BW (g) | 90±3.67 | | 93.73±2.42 | | 97.70±4.23 | | 0.36 |
| **S4** | BW (g) | 462.53±19.23 | **b** | 539.20±8.49 | **a** | 474.17±3.95 | **b** | **0.009** |
| | CF | 1.46±0.01 | **b** | 1.55±0.01 | **a** | 1.53±0.02 | **a** | **0.006** |
| | HSI | 1.64±0.08 | **b** | 1.32±0.04 | **a** | 1.42±0.05 | **ab** | **0.025** |

Mean values of three tanks are followed by SEM and the compact letter display of Tukey's post hoc test ($p < 0.05$, ANOVA followed by Tukey's post hoc test).
†BW (body weight), CF (condition factor), and HSI (hepatosomatic index).

**Table S3.** Read counts of RNA-seq samples after initial quality control, alignment, and quantification.

| Sample | Sex | Treatment | M Seqs[1] | M Aligned[2] | % Aligned[2] | M Assigned[3] | % Assigned[3] |
|---|---|---|---|---|---|---|---|
| **1Cp1** | M | 1C+ | 14.6 | 12 | 82.1% | 10.8 | 58.9% |
| **1Cp2** | M | 1C+ | 11.8 | 9.6 | 81.8% | 8.6 | 57.5% |
| **1Cp3** | M | 1C+ | 16.8 | 13.7 | 81.7% | 12.3 | 58.2% |
| **1Cp4** | F | 1C+ | 14.4 | 11.7 | 80.7% | 10.5 | 55.4% |
| **1Cp5** | F | 1C+ | 15.8 | 12.9 | 81.8% | 11.7 | 58.5% |
| **1Cp6** | M | 1C+ | 14.5 | 11.6 | 79.8% | 10.4 | 53.9% |
| **1Cp7** | F | 1C+ | 16.8 | 13.7 | 81.9% | 12.3 | 58.6% |
| **1Cp8** | F | 1C+ | 16.8 | 13.2 | 78.5% | 12 | 56.4% |
| **1Cp9** | F | 1C+ | 15.5 | 12.7 | 82.1% | 11.4 | 58.8% |
| **1Cpp1** | F | 1C++ | 14.5 | 11.9 | 82.3% | 10.8 | 60.3% |
| **1Cpp2** | M | 1C++ | 16.5 | 13.4 | 81.2% | 12.2 | 57.7% |
| **1Cpp3** | F | 1C++ | 16.6 | 13.4 | 80.8% | 12.2 | 57.1% |
| **1Cpp4** | F | 1C++ | 16.1 | 13.2 | 82.1% | 11.8 | 57.8% |
| **1Cpp5** | F | 1C++ | 16.6 | 13.3 | 80.2% | 11.9 | 54.8% |
| **1Cpp6** | M | 1C++ | 14.3 | 11.6 | 81.3% | 10.4 | 56.9% |
| **1Cpp7** | F | 1C++ | 14.5 | 11.5 | 79.3% | 10.4 | 53.0% |
| **1Cpp8** | M | 1C++ | 14.6 | 12 | 82.2% | 10.8 | 58.9% |
| **1Cpp9** | M | 1C++ | 16.2 | 13.2 | 81.8% | 12 | 58.5% |
| **Ctrl1** | F | Ctrl | 17.1 | 13.9 | 81.4% | 12.5 | 57.3% |
| **Ctrl2** | F | Ctrl | 12.4 | 10.3 | 82.8% | 9.2 | 60.3% |
| **Ctrl3** | M | Ctrl | 15.4 | 12.6 | 81.8% | 11.3 | 58.1% |
| **Ctrl4** | F | Ctrl | 18 | 14.9 | 82.7% | 13.4 | 59.8% |
| **Ctrl5** | M | Ctrl | 15.8 | 13.1 | 82.9% | 11.8 | 60.4% |
| **Ctrl6** | M | Ctrl | 18.3 | 14.6 | 79.6% | 13 | 52.2% |
| **Ctrl7** | F | Ctrl | 15.5 | 12.9 | 83.1% | 11.6 | 60.4% |
| **Ctrl8** | M | Ctrl | 17.9 | 14.7 | 82.2% | 13.3 | 59.2% |
| **Ctrl9** | F | Ctrl | 16.4 | 13.3 | 81.3% | 11.9 | 56.5% |

[1]Total read count after initial quality control by Trim Galore!.
[2]Count of aligned reads to the reference genome by STAR and the percentage of the aligned reads to the total reads.
[3]Count of the reads associated with known RNAs by featureCount with the percentage of the assigned reads among the total aligned sites, which include both unique and multiple aligned reads.

**Table S4.** Number of DEGs identified by three comparisons.

| Comparison | Control | # DEGs | # Down-regulated | # Up-regulated |
|---|---|---|---|---|
| **1C+ vs Ctrl** | Ctrl | 874 | 513 | 361 |
| **1C++ vs Ctrl** | Ctrl | 759 | 395 | 364 |
| **1C++ vs 1C+** | 1C+ | 20 | 10 | 10 |

**Table S5.** Enriched KEGG pathways for the genes in DEG C1 and DEG C2 clusters by ORA.

| Cluster | ID | Description | GeneRatio[1] | BgRatio[3] | p.adjust[3] | GSEA[4] |
|---|---|---|---|---|---|---|
| **DEG C1** | sasa01240 | Biosynthesis of cofactors | 17/213 | 201/7462 | 7.48E-03 | Y |
| | sasa00270 | Cysteine and methionine metabolism | 10/213 | 86/7462 | 1.03E-02 | Y |
| | sasa00220 | Arginine biosynthesis | 6/213 | 34/7462 | 1.54E-02 | N |
| | sasa01230 | Biosynthesis of amino acids | 11/213 | 127/7462 | 2.98E-02 | Y |
| | sasa00982 | Drug metabolism - cytochrome P450 | 6/213 | 42/7462 | 2.98E-02 | N |
| **DEG C2** | sasa00591 | Linoleic acid metabolism | 4/144 | 19/7462 | 2.15E-02 | Y |
| | sasa04141 | Protein processing in endoplasmic reticulum | 16/144 | 321/7462 | 2.15E-02 | Y |
| | sasa03060 | Protein export | 5/144 | 40/7462 | 3.15E-02 | Y |
| | sasa00980 | Metabolism of xenobiotics by cytochrome P450 | 5/144 | 43/7462 | 3.30E-02 | N |

[1,2,3]Output of the enrichKEGG function provided by the clusterProfiler package. GeneRatio: gene ratio, BgRatio: background ratio, p.adjust: adjusted p-value by the Benjamini-Hochberg procedure.
[4]Y: the pathway is also enriched by one of the GSEA results. N: the pathway is not enriched by GSEA.

**Table S6.** Enriched KEGG pathways for the C1+ vs Ctrl comparison by GSEA.

| ID | Description | setSize[1] | NES[2] | p.adjus[3] | Support[4] |
|---|---|---|---|---|---|
| **sasa03010** | Ribosome | 255 | 2.09E+00 | 5.33E-09 | GSEA |
| **sasa04141** | Protein processing in endoplasmic reticulum | 378 | 1.95E+00 | 5.33E-09 | ORA, GSEA |
| **sasa04110** | Cell cycle | 275 | -1.94E+00 | 5.33E-09 | GSEA |
| **sasa04115** | p53 signaling pathway | 145 | -1.82E+00 | 1.04E-04 | |
| **sasa04510** | Focal adhesion | 470 | 1.48E+00 | 1.51E-04 | |
| **sasa03060** | Protein export | 44 | 2.10E+00 | 1.70E-04 | ORA |
| **sasa04216** | Ferroptosis | 108 | -1.84E+00 | 1.84E-04 | |
| **sasa04260** | Cardiac muscle contraction | 191 | 1.72E+00 | 1.84E-04 | |
| **sasa04068** | FoxO signaling pathway | 331 | -1.58E+00 | 2.24E-04 | |
| **sasa04218** | Cellular senescence | 373 | -1.57E+00 | 2.24E-04 | |
| **sasa04914** | Progesterone-mediated oocyte maturation | 203 | -1.67E+00 | 2.58E-04 | |
| **sasa00190** | Oxidative phosphorylation | 245 | 1.61E+00 | 3.20E-04 | |
| **sasa00100** | Steroid biosynthesis | 31 | -2.02E+00 | 5.46E-04 | GSEA |
| **sasa01230** | Biosynthesis of amino acids | 167 | -1.64E+00 | 7.34E-04 | ORA |
| **sasa00240** | Pyrimidine metabolism | 103 | -1.74E+00 | 7.68E-04 | |

[1,2,3]Output of the gseKEGG function provided by the clusterProfiler package. setSize: the number of genes that contributed for enrichment, NES: normalized enrichment score that indicates up-regulation (positive) or down-regulation (negative), p.adjust: adjusted p-value by the Benjamini-Hochberg procedure.
[4]ORA: the pathway is also enriched by ORA. GSEA: the pathway is also enriched by at least one of the other GSEA results.

**Table S7.** Enriched KEGG pathways for the C1++ vs Ctrl comparison by GSEA.

| ID | Description | setSize[1] | NES[2] | p.adjust[3] | Support[4] |
|---|---|---|---|---|---|
| **sasa03010** | Ribosome | 255 | 2.63E+00 | 1.60E-08 | GSEA |
| **sasa00970** | Aminoacyl-tRNA biosynthesis | 62 | -2.26E+00 | 4.86E-07 | GSEA |
| **sasa04110** | Cell cycle | 275 | -1.83E+00 | 1.52E-06 | GSEA |
| **sasa00190** | Oxidative phosphorylation | 247 | 1.79E+00 | 6.07E-06 | |
| **sasa03030** | DNA replication | 55 | -2.01E+00 | 1.47E-04 | |
| **sasa01230** | Biosynthesis of amino acids | 164 | -1.81E+00 | 1.47E-04 | |
| **sasa00270** | Cysteine and methionine metabolism | 107 | -1.85E+00 | 1.87E-04 | |
| **sasa00100** | Steroid biosynthesis | 32 | -2.04E+00 | 3.83E-04 | GSEA |
| **sasa01232** | Nucleotide metabolism | 164 | -1.72E+00 | 5.59E-04 | |
| **sasa01240** | Biosynthesis of cofactors | 260 | -1.62E+00 | 5.59E-04 | ORA |

[1,2,3]Output of the gseKEGG function provided by the clusterProfiler package. setSize: the number of genes that contributed for enrichment, NES: normalized enrichment score that indicates up-regulation (positive) or down-regulation (negative), p.adjust: adjusted p-value by the Benjamini-Hochberg procedure.
[4]ORA: the pathway is also enriched by ORA. GSEA: the pathway is also enriched by at least one of the other GSEA results.

**Table S8.** Enriched KEGG pathways for the C1+ vs Ctrl comparison by GSEA.

| ID | Description | setSize[1] | NES[2] | p.adjust[3] | Support[4] |
|---|---|---|---|---|---|
| **sasa03010** | Ribosome | 257 | 1.88E+00 | 3.43E-07 | GSEA |
| **sasa00970** | Aminoacyl-tRNA biosynthesis | 61 | -2.24E+00 | 4.56E-07 | GSEA |
| **sasa03015** | mRNA surveillance pathway | 179 | -1.66E+00 | 2.76E-04 | |
| **sasa04141** | Protein processing in endoplasmic reticulum | 375 | -1.54E+00 | 4.17E-04 | ORA,GSEA |
| **sasa03013** | Nucleocytoplasmic transport | 209 | -1.67E+00 | 4.33E-04 | |

[1,2,3]Output of the gseKEGG function provided by the clusterProfiler package. setSize: the number of genes that contributed for enrichment, NES: normalized enrichment score that indicates up-regulation (positive) or down-regulation (negative), p.adjust: adjusted p-value by the Benjamini-Hochberg procedure.
[4]ORA: the pathway is also enriched by ORA. GSEA: the pathway is also enriched by at least one of the other GSEA results.

**Table S9.** Read counts of RRBS samples after initial quality control and alignment percentage.

| Sample Name | Sex | Treatment | M Seqs[1] | % Aligned[2] |
|---|---|---|---|---|
| **1Cp1** | F | 1C+ | 67.4 | 47.5% |
| **1Cp2** | F | 1C+ | 44.7 | 47.5% |
| **1Cp3** | M | 1C+ | 74.2 | 47.1% |
| **1Cp4** | F | 1C+ | 46.7 | 47.1% |
| **1Cp5** | M | 1C+ | 51.6 | 47.8% |
| **1Cp6** | M | 1C+ | 49.1 | 47.1% |
| **1Cp7** | F | 1C+ | 42.3 | 46.7% |
| **1Cp8** | M | 1C+ | 66.3 | 47.0% |
| **1Cp9** | F | 1C+ | 58.6 | 46.8% |
| **1Cpp1** | M | 1C++ | 61.8 | 48.3% |
| **1Cpp2** | M | 1C++ | 61.6 | 49.0% |
| **1Cpp3** | M | 1C++ | 54.4 | 48.1% |
| **1Cpp4** | F | 1C++ | 66.3 | 47.5% |
| **1Cpp5** | F | 1C++ | 47.8 | 48.4% |
| **1Cpp6** | M | 1C++ | 36.5 | 47.7% |
| **1Cpp7** | F | 1C++ | 59.6 | 46.9% |
| **1Cpp8** | F | 1C++ | 50.7 | 46.4% |
| **1Cpp9** | F | 1C++ | 59.7 | 47.2% |
| **Ctrl1** | F | Ctrl | 63 | 48.0% |
| **Ctrl2** | M | Ctrl | 59 | 47.8% |
| **Ctrl3** | F | Ctrl | 65.9 | 47.9% |
| **Ctrl4** | F | Ctrl | 42.4 | 47.0% |
| **Ctrl5** | F | Ctrl | 60.3 | 46.7% |
| **Ctrl6** | M | Ctrl | 50.5 | 46.7% |
| **Ctrl7** | F | Ctrl | 46.4 | 47.4% |
| **Ctrl8** | M | Ctrl | 55 | 47.9% |
| **Ctrl9** | M | Ctrl | 45.4 | 47.6% |

[1]Total read count after initial quality control by Trim Galore!.
[2] Percentage of the aligned reads to the reference genome by Bismark.

**Table S10.** Comparisons of two methylation rate distributions in different regions by KS tests.

| Region | Size | X | Y | Alternative | p-value | Significance† |
|---|---|---|---|---|---|---|
| **All mapped CpGs** | 157 201 | 1C+ | Ctrl | **less** | **0** | * |
| | | | | greater | 0.81 | |
| | | 1C++ | Ctrl | **less** | **0** | * |
| | | | | greater | 0.93 | |
| | | 1C+ | 1C++ | **less** | **0** | * |
| | | | | greater | 0.85 | |
| **GB** | 87 215 | 1C+ | Ctrl | **less** | **0** | * |
| | | | | greater | 0.83 | |
| | | 1C++ | Ctrl | **less** | **0** | * |
| | | | | greater | 0.91 | |
| | | 1C+ | 1C++ | **less** | **8.88e-16** | * |
| | | | | greater | 0.85 | |
| **P** | 3 091 | 1C+ | Ctrl | **less** | **0.04** | * |
| | | | | greater | 0.81 | |
| | | 1C++ | Ctrl | **less** | **0.02** | * |
| | | | | greater | 0.9 | |
| | | 1C+ | 1C++ | less | 0.72 | |
| | | | | greater | 0.47 | |
| **RS** | 48 148 | 1C+ | Ctrl | **less** | **0** | * |
| | | | | greater | 0.88 | |
| | | 1C++ | Ctrl | **less** | **0** | * |
| | | | | greater | 0.84 | |
| | | 1C+ | 1C++ | **less** | **9.55e-10** | * |
| | | | | greater | 0.93 | |
| **IGR** | 47 748 | 1C+ | Ctrl | **less** | **0** | * |
| | | | | greater | 0.93 | |
| | | 1C++ | Ctrl | **less** | **0** | * |
| | | | | greater | 0.98 | |
| | | 1C+ | 1C++ | **less** | **0** | * |
| | | | | greater | 0.94 | |

†'*' indicates that KS test result is statistically significant with p-value < 0.05.

**Table S11.** Number of DMCs identified by three comparisons in four different regions.

| Comparison | Region | #Mapped CpGs | #DMCs | (%)[1] | #DMCs (hypo)[2] | #DMCs (hyper)[3] |
|---|---|---|---|---|---|---|
| **1C+ vs Ctrl** | GB | 107366 | 3061 | 2.9% | 933 | 2128 |
| | P | 4189 | 154 | 3.7% | 49 | 105 |
| | RS | 74773 | 2390 | 3.2% | 793 | 1597 |
| | IGR | 67078 | 2112 | 3.1% | 636 | 1476 |
| **1C++ vs Ctrl** | GB | 108055 | 2969 | 2.7% | 978 | 1991 |
| | P | 4110 | 131 | 3.2% | 58 | 73 |
| | RS | 75504 | 2145 | 2.8% | 729 | 1416 |
| | IGR | 67018 | 2044 | 3% | 652 | 1392 |
| **1C++ vs 1C+** | GB | 105087 | 2488 | 2.4% | 1359 | 1129 |
| | P | 4103 | 134 | 3.3% | 84 | 50 |
| | RS | 74515 | 1973 | 2.6% | 1048 | 925 |
| | IGR | 65076 | 1766 | 2.7% | 967 | 799 |

[1]Percentage of the DMC count calculated by (#DMCs)/(#Mapped CpGs) * 100.
[2,3]Number of hypo-methylated and hyper-methylated DMCs receptively.

**Table S12.** List of genes that have multiple DMCs in the promoter (P) regions.

| Comparison | Gene ID[1] | Gene symbol[2] | Gene name | #DMCs[3] |
|---|---|---|---|---|
| **1C+ vs Ctrl** | 100195955 | rn182 | RING finger protein 182 | 6 (0/6) |
| | 106572013 | LOC106572013 | uncharacterized LOC106572013 | 4 (0/4) |
| | 106602923 | LOC106602923 | serine/threonine/tyrosine-interacting protein A-like | 3 (0/3) |
| **1C++ vs Ctrl** | 100195955 | rn182 | RING finger protein 182 | 4 (0/4) |
| | 100196228 | yars | tyrosyl-tRNA synthetase | 4 (4/0) |
| | 106605303 | LOC106605303 | lysyl oxidase homolog 3-like | 3 (0/3) |
| **1C++ vs 1C+** | 100195786 | nca11 | Neural cell adhesion molecule 1-A | 3 (3/0) |
| | 106572013 | LOC106572013 | uncharacterized LOC106572013 | 4 (4/0) |
| | 106588671 | LOC106588671 | DNA topoisomerase 2-beta-like | 2 (0/2) |

[1,2]Gene ID and gene symbol from NCBI,
[3]Number of DMCs identified in the promoter region with (hypo-methylated/hyper-methylated).

**Table S13.** List of genes that have multiple DMCs in the RS (5K) regions.

| Select[1] | Comparison | Gene ID[2] | Gene symbol[3] | Gene name | #DMCs[4] |
|---|---|---|---|---|---|
| **Direct** | 1C+ vs Ctrl | 106574560 | LOC106574560 | cysteine/serine-rich nuclear protein 3-like | 5 (5/0) |
| | | 106577634 | LOC106577634 | uncharacterized LOC106577634 | 6 (0/6) |
| | | 106586627 | LOC106586627 | vang-like protein 1 | 5 (0/5) |
| | 1C++ vs Ctrl | 106577634 | LOC106577634 | uncharacterized LOC106577634 | 4 (0/4) |
| | | 106604632 | LOC106604632 | WD40 repeat-containing protein SMU1 | 3 (0/3) |
| | | 106610962 | LOC106610962 | transcription factor IIIB 90 kDa subunit-like | 4 (4/0) |
| | 1C++ vs 1C+ | 106586604 | tbx15 | T-box 15 | 3 (1/2) |
| | | 106591533 | btg4 | B-cell translocation gene 4 | 8 (0/8) |
| | | 106609646 | LOC106609646 | hepatocyte growth factor-like | 4 (0/4) |
| **In-direct** | 1C+ vs Ctrl | 106591533 | btg4 | B-cell translocation gene 4 | 2 (2/0) |
| | | 106604632 | LOC106604632 | WD40 repeat-containing protein SMU1 | 1 (0/1) |
| | | 106610962 | LOC106610962 | transcription factor IIIB 90 kDa subunit-like | 2 (2/0) |
| | 1C++ vs Ctrl | 106609646 | LOC106609646 | hepatocyte growth factor-like | 1 (0/1) |
| | 1C++ vs 1C+ | 106574560 | LOC106574560 | cysteine/serine-rich nuclear protein 3-like | 2 (0/2) |
| | | 106604632 | LOC106604632 | WD40 repeat-containing protein SMU1 | 2 (0/2) |

[1]"Direct" and "In-direct" show how the genes are identified. Direct selection is liked to top 3 genes when the genes are sorted by the number of DMCs by descendent order within one of the three comparisons. In-direct selection is simply added when a gene is identified by the "direct" selection and has at least one DMC in the RS (5K) region within other comparisons.

[2,3]Gene ID and gene symbol from NCBI,

[4]Number of DMCs identified in the RS (5K) region with (hypo-methylated/hyper-methylated).

**Table S14.** List of genes that have multiple DMCs in the GB (exon) regions.

| Select[1] | Comparison | Gene ID[2] | Gene symbol[3] | Gene name | #DMCs[4] |
|---|---|---|---|---|---|
| **Direct** | 1C+ vs Ctrl | 106571647 | st8sia4 | ST8 alpha-N-acetyl-neuraminide alpha-2,8-sialyltransferase 4 | 5 (5/0) |
| | | 106574559 | LOC106574559 | xin actin-binding repeat-containing protein 2-like | 5 (5/0) |
| | | 106577636 | LOC106577636 | RING finger protein 186-like | 6 (0/6) |
| | 1C++ vs Ctrl | 106577636 | LOC106577636 | RING finger protein 186-like | 4 (0/4) |
| | | 106590562 | LOC106590562 | complement C3-like | 4 (4/0) |
| | | 106608642 | LOC106608642 | CD276 antigen-like | 5 (5/0) |
| | 1C++ vs 1C+ | 106564966 | LOC106564966 | beta-1,3-galactosyl-O-glycosyl-glycoprotein beta-1,6-N-acetylglucosaminyltransferase 3-like | 4 (0/4) |
| | | 106571647 | st8sia4 | ST8 alpha-N-acetyl-neuraminide alpha-2,8-sialyltransferase 4 | 6 (0/6) |
| | | 106599566 | LOC106599566 | protein LBH-like | 4 (4/0) |
| **In-direct** | 1C+ vs Ctrl | 106590562 | LOC106590562 | complement C3-like | 4 (4/0) |
| | | 106599566 | LOC106599566 | protein LBH-like | 4 (0/4) |
| | 1C++ vs Ctrl | 106564966 | LOC106564966 | beta-1,3-galactosyl-O-glycosyl-glycoprotein beta-1,6-N-acetylglucosaminyltransferase 3-like | 3 (0/3) |
| | | 106599566 | LOC106599566 | protein LBH-like | 2 (2/0) |
| | 1C++ vs 1C+ | 106608642 | LOC106608642 | CD276 antigen-like | 4 (4/0) |

[1]"Direct" and "In-direct" show how the genes are identified. Direct selection is liked to top 3 genes when the genes are sorted by the number of DMCs by descendent order within one of the three comparisons. In-direct selection is simply added when a gene is identified by the "direct" selection and has at least one DMC in the GB (exon) region within other comparisons.

[2,3]Gene ID and gene symbol from NCBI,

[4]Number of DMCs identified in the GB (exon) region with (hypo-methylated/hyper-methylated).

**Table S15.** List of genes that are DEGs and contain DMRs around their TSSs.

| Comp | Region | Gene ID[1] | Gene symbol[2] | Gene name | Dist[3] | LFC[4] | Mdiff[5] |
|---|---|---|---|---|---|---|---|
| **1C+ vs Ctrl** | GB (exon) | 106560487 | angptl3 | angiopoietin-like 3 | 553 | -1.53 | 23.80 |
| | | 106600884 | LOC106600884 | 3-mercaptopyruvate sulfurtransferase-like | 971 | 0.34 | 19.57 |
| | P | 106593742 | LOC106593742 | group 10 secretory phospholipase A2-like | -636 | 1.08 | 15.19 |
| | | 106604118 | LOC106604118 | PRELI domain-containing protein 1, mitochondrial-like | -829 | 0.31 | 24.21 |
| | | 106603181 | LOC106603181 | organic solute transporter subunit alpha-like | -869 | -0.62 | -16.06 |
| | | 106584206 | LOC106584206 | FYN-binding protein 1 | -451 | 1.39 | -16.21 |
| **1C++ vs Ctrl** | GB (exon) | 106580755 | LOC106580755 | cytochrome P450 2M1-like | 280 | -8.41 | 15.14 |
| | | 100380841 | slc3a1 | solute carrier family 3 (amino acid transporter heavy chain), member 1 | 157 | -0.99 | 17.94 |
| | P | 106604118 | LOC106604118 | PRELI domain-containing protein 1, mitochondrial-like | -829 | 0.34 | 18.16 |
| | | 106611251 | LOC106611251 | HHIP-like protein 1 | -183 | -0.93 | -16.79 |

[1,2]Gene ID and gene symbol from NCBI.
[3]Distance from TSS.
[4]Log fold changes of DEGs.
[5]Methylation differences of DMRs.

**Table S16.** List of DEGs from three comparisons: 1C+ vs Ctrl, 1C++ vs Ctrl, and 1C++ vs 1C+.

**File:** 01_degs_total.xlsx (provided in Excel format)

**Sheets:** 1C+ vs Ctrl, 1C++ vs Ctrl, 1C++ vs 1C+

**Fields:**

| | |
|---|---|
| gene_id | Entrez Gene ID from NCBI |
| lfc | Log fold change produced by DESeq2 |
| padj | Adjusted p-value produced by DESeq2 |
| type | RNA type |
| gene_symbol | Gene symbol from NCBI |
| gene_name | Gene name from NCBI |
| orgdb_gene_symbol | Gene symbol from BioConductor OrgDB |
| orgdb_gene_name | Gene name from BioConductor OrgDB |
| orgdb_alias | Gene alias from BioConductor OrgDB |

**Table S17.** List of DEGs from three comparisons: 1C+ vs Ctrl, 1C++ vs Ctrl, and 1C++ vs 1C+.

**File:** 02_degs_stringent.xlsx (provided in Excel format)

**Sheets:** 1C+ vs Ctrl, 1C++ vs Ctrl

**Fields:**

| | |
|---|---|
| gene_id | Entrez Gene ID from NCBI |
| lfc | Log fold change produced by DESeq2 |
| padj | Adjusted p-value produced by DESeq2 |
| gene_symbol | Gene symbol from NCBI |
| gene_name | Gene name from NCBI |

**Table S18.** List of DEGs from three comparisons: 1C+ vs Ctrl, 1C++ vs Ctrl, and 1C++ vs 1C+.

**File:** 03_degs_stringent_relaxed.xlsx (provided in Excel format)

**Sheets:** 1C+ vs Ctrl, 1C++ vs Ctrl, 1C++ vs 1C+

**Fields:**

| | |
|---|---|
| gene_id | Entrez Gene ID from NCBI |
| lfc | Log fold change produced by DESeq2 |
| padj | Adjusted p-value produced by DESeq2 |
| gene_symbol | Gene symbol from NCBI |
| gene_name | Gene name from NCBI |

**Table S19.** List of DEG clusters identified by DBSCAN.

**File:** 04_dbscan_clusters.xlsx (provided in Excel format)

**Sheet:** DBSCAN

**Fields:**

| | |
|---|---|
| gene_id | Entrez Gene ID from NCBI |
| cluster | Cluster name identified by DBSCAN |
| type | RNA type |
| gene_symbol | Gene symbol from NCBI |
| gene_name | Gene name from NCBI |
| orgdb_gene_symbol | Gene symbol from BioConductor OrgDB |
| orgdb_gene_name | Gene name from BioConductor OrgDB |
| orgdb_alias | Gene alias from BioConductor OrgDB |

**Table S20.** List of enriched KEGG pathways identified by ORA and GSEA.

**File:** 05_kegg_rna.xlsx (provided in Excel format)

**Sheets:** ORA DEG C1, ORA DEG C2

**Fields:**

| | |
|---|---|
| ID | KEGG ID |
| Description | KEGG pathway |
| GeneRatio | Gene ratio used in ORA calculation |
| BgRatio | Back ground ratio used in ORA calculation |
| pvalue | p-value calculated by clusterProfiler |
| p.adjust | Adjusted p-value calculated by clusterProfiler |
| qvalue | Q-value calculated by clusterProfiler |
| geneID | Gene IDs of DEGs involvoed in the corresponding KEGG pathway |
| Count | Count of DEGs involvoed in the corresponding KEGG pathway |

**Sheets:** GSEA 1C+, GSEA 1C++, GSEA 1C++ vs 1C+

**Fields:**

| | |
|---|---|
| Comparison | Comparison result used for GSEA analysis |
| ID | KEGG ID |
| Description | KEGG pathway |
| setSize | The size of gene set used in GSEA calculation |
| enrichmentScore | Enrichment score calcualted by clusterProfiler |
| NES | Normalised enrichment score calcualted by clusterProfiler |
| pvalue | p-value calculated by clusterProfiler |
| p.adjust | Adjusted p-value calculated by clusterProfiler |
| qvalue | Q-value calculated by clusterProfiler |
| rank | Rank calculated by clusterProfiler |
| leading_edge | Leading edge analysis performed by clusterProfiler |
| core_enrichment | Gene names that contibuted to enrichemnt |

**Table S21.** List of DMCs from three comparisons: 1C+ vs Ctrl, 1C++ vs Ctrl, and 1C++ vs 1C+.

**File:** 06_dmc_md15.xlsx (provided in Excel format)

**Sheets:** 1C+ vs Ctrl, 1C++ vs Ctrl, 1C++ vs 1C+

**Fields:**

| | |
|---|---|
| chrom | Chromosome of DMC |
| start | Start position of DMC |
| end | end position of DMC |
| strand | Strand of DMC |
| pvalue | p-value calculated by methylKit |
| qvalue | Q-value calculated by methylKit |
| meth.diff | Difference of methylation rate (%) |
| region | Region |
| gene_id | Gene ID from NCBI |
| gene_symbol | Gene symbol from NCBI |
| gene_name | Gene name from NCBI |
| refseq | Refseq ID |
| dist | Distance from TSS |

**Table S22.** List of DMRs from three comparisons: 1C+ vs Ctrl, 1C++ vs Ctrl, and 1C++ vs 1C+.

**File:** 07_dmr_md15.xlsx (provided in Excel format)

**Sheets:** 1C+ vs Ctrl, 1C++ vs Ctrl, 1C++ vs 1C+

**Fields:**

| | |
|---|---|
| chrom | Chromosome of DMR |
| start | Start position of DMR |
| end | end position of DMR |
| strand | Strand of DMR |
| pvalue | p-value calculated by methylKit |
| qvalue | Q-value calculated by methylKit |
| meth.diff | Difference of methylation rate (%) |
| region | Region |
| gene_id | Gene ID from NCBI |
| gene_symbol | Gene symbol from NCBI |
| gene_name | Gene name from NCBI |
| refseq | Refseq ID |
| dist | Distance from TSS |

### Supplementary figure

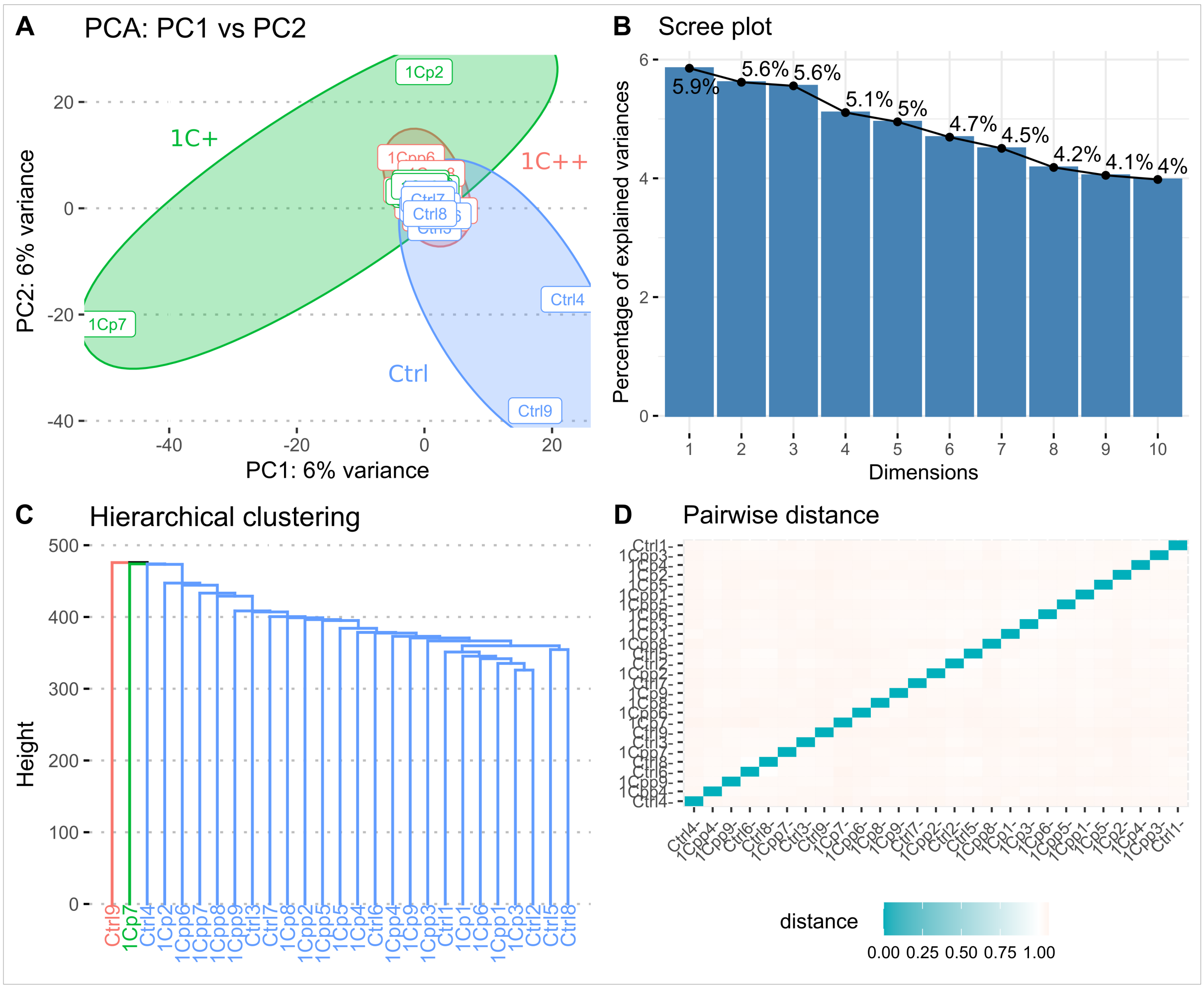


**Fig S1.** Clustering analysis on the methylation rates of mapped CpG.
Four different plots show the results of clustering analysis performed on the CpG sites with top 50% high variances. **(A)** PCA (principal component analysis) plot showing PC1 and PC2 components with three ellipses representing 1C+ (green), 1C++ (red), and Ctrl (blue). **(B)** Scree plot showing the percentage of explained variances for 10 principal components (PC1 ~ PC10). **(C)** The dendrogram showing the result of hierarchical clustering analysis. **(D)** Heatmap showing pairwise distances as the results of pairwise correlation analysis.
